# Using ‘sentinel’ plants to improve early detection of invasive plant pathogens

**DOI:** 10.1101/2022.09.01.506171

**Authors:** Francesca A. Lovell-Read, Stephen R. Parnell, Nik J. Cunniffe, Robin N. Thompson

## Abstract

Infectious diseases of plants present an ongoing and increasing threat to international biosecurity, with wide-ranging implications. An important challenge in plant disease management is achieving early detection of invading pathogens in new locations, which requires effective surveillance through the implementation of appropriate monitoring programs. However, when monitoring relies on visual inspection as a means of detection, surveillance is often hindered by a long incubation period (delay from infection to symptom onset) during which plants may be infectious but not displaying visible symptoms. ‘Sentinel’ plants – alternative susceptible host species that display visible symptoms of infection more rapidly – could be introduced to at-risk populations and included in monitoring programs to act as early warning beacons for infection. However, while sentinel hosts exhibit faster disease progression and so allow pathogens to be detected earlier, this often comes at a cost: faster disease progression typically promotes earlier onward transmission. Here, we construct a computational model of pathogen transmission to explore this trade-off and investigate how including sentinel plants in monitoring programmes could facilitate earlier detection of invasive plant pathogens. Using *Xylella fastidiosa* infection in *Olea europaea* (European olive) as a current high profile case study, for which *Catharanthus roseus* (Madagascan periwinkle) is a candidate sentinel host, we apply a Bayesian optimisation algorithm to determine the optimal number of sentinel hosts to introduce for a given sampling effort, as well as the optimal division of limited surveillance resources between crop and sentinel plants. Our results demonstrate that including sentinel plants in monitoring programmes can reduce the expected prevalence of infection upon outbreak detection substantially, increasing the feasibility of local outbreak containment.

## 1. Introduction

Infectious disease outbreaks in plant populations are responsible for devastating economic, environmental and societal consequences [1–8]. The global trade in live plant species means that the spread of invasive plant pathogens poses an ever-increasing threat to international biosecurity [9, 10]. Developing efficient and cost-effective methods for surveillance and control of invasive plant pathogens is therefore a vital area of current research [11–16].

Mathematical modelling is increasingly used to guide surveillance and intervention strategies for plant pathogens [11, 14, 17–19], helping policy-makers understand how to direct limited resources for control to reduce transmission [20–25]. Multiple studies of different pathogens have focused on the question of how to optimise control measures when a pathogen is known to be in a particular host landscape (‘reactive’ control). For example, using citrus canker (a bacterial disease of citrus plants) in Florida as a case study, Cunniffe *et al*. [26] showed how roguing (removal of confirmed infected plants) can be extended to removal of all plants in the proximity of a confirmed infected host. They demonstrated a modelling approach for identifying the optimal removal radius around confirmed infected hosts, balancing the reduction in the risk of further spread with the cost of destroying potentially healthy plants. Similarly, White *et al*. [27] investigated the efficacy of ‘buffer zones’ for intensive surveillance on the border of a region of known infection, and Adrakey *et al*. [28] developed a Bayesian approach for prioritising the removal of infected hosts based on the infection risk they pose to other plants.

While reactive control has been well-studied, a key additional challenge is designing surveillance methods specifically to achieve early outbreak detection [29]. This increases the chance of eradicating the pathogen from the landscape before it becomes widespread (‘preventive’ control). Parnell *et al*. [30, 31] used probabilistic techniques to derive a simple ‘rule of thumb’ relating the expected prevalence of infection upon discovery to the sampling effort. This approach was extended by Mastin *et al*. [32] to a setting in which pathogen transmission via vectors is modelled explicitly, and used to investigate how to divide sampling resources optimally between hosts and vectors. Alonso-Chavez *et al*. [33] also applied this technique to explore the use of within-nursery surveillance for reducing the risk of growers selling infected plant material.

For many plant pathogens, a major obstacle to achieving early detection is a long incubation period (the time between initial infection and symptom onset; this has also been referred to as the cryptic, asymptomatic or presymptomatic period) [18, 33–38]. If transmission by infected hosts can occur prior to symptom onset, monitoring programs that rely upon visual inspection of potential hosts for signs of infection (as are standard across plant health [39, 40]) may fail to identify the presence of a pathogen before widespread transmission has occurred. Indeed, Alonso-Chavez *et al*. [33] showed that if presymptomatic transmission occurs and the pathogen is discovered early in an outbreak, the expected prevalence of infection in the population at the time of discovery increases exponentially with the duration of the incubation period.

The incubation period of a pathogen may, however, vary substantially between different host species, or between hosts of different ages [35–37, 41, 42]. This suggests that in some cases there may be the potential to use alternative hosts with relatively short incubation periods as ‘sentinels’ to detect new outbreaks. In this context, we refer to a ‘sentinel’ as a susceptible plant species specifically chosen to have a short incubation period, which is placed amongst the crop plants and regularly monitored for visible signs of infection (we note that, in the literature, the term ‘sentinel’ has alternatively been used to refer to plant species that are grown outside of their natural habitat and monitored to assess whether pests native to the new location pose a risk to that species, which is distinct from the context of this study [43]).

The rapid onset of visible symptoms in sentinel plants could result in earlier detection of the pathogen in the population. If this leads to a reduced prevalence of infection upon discovery, this could reduce the cost of reactive control as well as increasing the probability of successful pathogen eradication following detection. However, sentinels may also have drawbacks. The increased rate of symptom development in sentinel plants could result in more rapid onward transmission, which may counteract the positive effect of early detection and therefore lead to an increased discovery prevalence. Therefore, research is needed to understand this trade-off and infer the conditions under which sentinel plants are likely to be beneficial for reducing the discovery prevalence. If sentinel plants are deployed, one key consideration is how many sentinel plants should be added to the population to provide sufficient opportunity for early detection whilst limiting the concurrent increase in the transmission rate. Another important question is how to divide limited sampling resources optimally between crop and sentinel plants. Is it better to sample preferentially from the available sentinel population, or should a mixture of crop and sentinel plants be inspected?

Although using sentinel plants (as defined in the context of this study) to facilitate early detection of invasive pathogens has been suggested as a possibility previously [43, 44], the question of how to design effective surveillance strategies using sentinel hosts targeted at a specific pathogen in a given region has not yet been addressed [43]. Here, we explore the potential for sentinel plants to aid early outbreak detection, using a plant pathogen of significant current importance as a case study (*Xylella fastidiosa* – see below). We construct a stochastic compartmental model of pathogen transmission that includes two different host types (crops and sentinels), and consider monitoring programs in which a fixed number of plants are selected at random and inspected for visible disease symptoms at regular intervals. A surveillance strategy is defined by: i) the number of sentinel plants added to the population; ii) the number of crops and sentinels to be examined in each sampling round, and; iii) the time interval between successive sampling rounds. For a given surveillance strategy, we use model simulations to calculate the expected detection prevalence (EDP) of the pathogen in the crop population at the time of discovery. We investigate the conditions under which including sentinel plants in a surveillance programme allows us to attain a lower EDP than standard monitoring (i.e., the analogous surveillance strategy but without any sentinel plants). For a given choice of sample size and sample interval, we apply a Bayesian optimisation algorithm to determine the minimum attainable EDP and the surveillance strategy for which this is achieved.

We demonstrate that including sentinel plants in a surveillance programme has the potential to reduce the EDP compared to a standard monitoring programme of equivalent sampling effort. Sentinel plants are particularly beneficial when limited resources are available for plant disease surveillance. We show that both the total number of sentinels deployed and the division of the sample between crop and sentinel plants are crucial in determining the effectiveness of a surveillance strategy. It can be preferable to sample a mixture of both sentinel and crop plants, rather than exclusively sampling sentinels. Overall, our results demonstrate that sentinel plants are a useful tool to improve early detection monitoring, and encourage further research to identify the range of host-pathogen systems for which sentinel plants can reduce the damage caused by invading plant pathogens.

### Case study: *Xylella fastidiosa* infection in *Olea europaea* (European olive)

An important example of a pathogen for which the development of effective surveillance strategies is currently critical is *Xylella fastidiosa* [45], a vector-borne bacterial pathogen first isolated from infected grapevines in 1978 [46]. It is pathogenic to over 600 host plant species [47], including economically important crops such as grapevines, almonds, olives, citrus and coffee [48, 49].

Depending on the host species and specific bacterial strain, symptoms of *X. fastidiosa* infection include leaf tissue necrosis (leaf scorch), stunted growth, decrease in fruit production, dieback and eventual death [36, 48, 49]. Outbreaks of *X. fastidiosa* in commercial crops therefore have substantial negative economic effects [22, 50, 51].

Of particular current concern is a strain of *X. fastidiosa*, subspecies *pauca*, known as CoDiRO (Complesso del Disseccamento Rapido dell’Olivo - loosely, ‘rapid drying disease of olive trees’), which was discovered in the Apulia region of south-east Italy in 2013 and subsequently identified as the causative agent of Olive Quick Decline Syndrome (OQDS) in that region [27, 52]. CoDiRO spreads rapidly, is difficult to contain, and results in significant crop loss, threatening olive farming throughout Europe [22, 35, 48, 53–55]. Recent projections indicate that the economic impact on olive farming in Italy, Greece and Spain alone could exceed €24 billion over the next 50 years if feasible control strategies are not devised [22]. However, the wide range of possible hosts for *X. fastidiosa* means that this pathogen poses a risk to European agriculture on an even broader scale [35, 54–56].

Despite ongoing research, there is currently no effective treatment for OQDS, or *X. fastidiosa* infection more generally [48, 49]. Control methods therefore mainly consist of roguing infected plants and removing healthy plants in their vicinity, and reducing vector transmission using insecticides [27]. However, these interventions are costly, and must be swift in order to be effective - once *X. fastidiosa* becomes established in a region, eradication becomes unfeasible [26, 55, 57]. Devising appropriate monitoring programmes to facilitate early detection is therefore critical to the success of containment strategies [35].

*X. fastidiosa* is a prime example of a pathogen for which the incubation period can provide a major obstacle to achieving early detection. *X. fastidiosa* subsp. *pauca*, the causal agent of OQDS, has a long incubation period in European olive (*Olea europaea*) with a mean duration of around 15 months [35], and transmission by infected hosts can occur prior to symptom onset [36]. Since surveillance strategies for OQDS typically rely upon visual inspection of potential hosts as a first line of defence [40, 48], presymptomatic transmission significantly limits the efficacy of current infection monitoring programs (although molecular tests are able to detect *X. fastidiosa* infection before symptom onset [58, 59], the costs of large-scale asymptomatic sampling are prohibitive [40]). Despite its long incubation period in *Olea europaea*, there is substantial variation in the incubation period of *X. fastidiosa* across its large host range, depending on factors such as plant species and age, pathogen subspecies, and climatic conditions [35, 36]. For this reason, the use of sentinel plants for surveillance of *X. fastidiosa* is a clear possibility, and has been identified as a key research area by the G20 Meetings of Agricultural Chief Scientists [36].

Here, we consider the candidate sentinel plant species *Catharanthus roseus*, a herbaceous flowering plant commonly known as Madagascan periwinkle. *C. roseus* is a known host of *X. fastidiosa* subsp. *pauca*, with a mean time from infection to symptom onset of around seven weeks (a factor of nine times shorter than *X. fastidiosa* in *O. europaea*) [35]. Although native to Madagascar, *C. roseus* has a wide geographical distribution and is already found across Italy and the rest of Europe, making it an ecologically suitable choice as a sentinel host [60]. Additionally, the small size of *C. roseus* plants allows them to be intercropped in olive groves.

Although here we use *X. fastidiosa* infection in *O. europaea* as an important case study, our modelling framework is intended to be general and extensible. It may also be applied to investigate the use of sentinels against other invasive pathogens for which asymptomatic infection hinders existing monitoring approaches.

## 2. Methods

### 2.1 Transmission model

We considered a compartmental model of pathogen transmission in which plants are classified as ‘Healthy’ (*H*), ‘Undetectable’ (*U*) or ‘Detectable’ (*D*). The model includes two host types – crop plants (denoted by subscript *C*) and sentinel plants (denoted by subscript *S*). ‘Healthy’ crops (*H*_*C*_) and sentinels (*H*_*S*_) are uninfected plants that are susceptible to infection. ‘Undetectable’ crops (*U*_*C*_) and sentinels (*U*_*S*_) are plants that have been infected (and are infectious) but are not currently displaying visual symptoms. Once visual symptoms develop, plants progress into the ‘Detectable’ compartment (*D*_*C*_ or *D*_*S*_ for crops and sentinels, respectively). A schematic illustrating the different compartments for both crop and sentinel plants is shown in Figure 1A.

**Fig 1.**
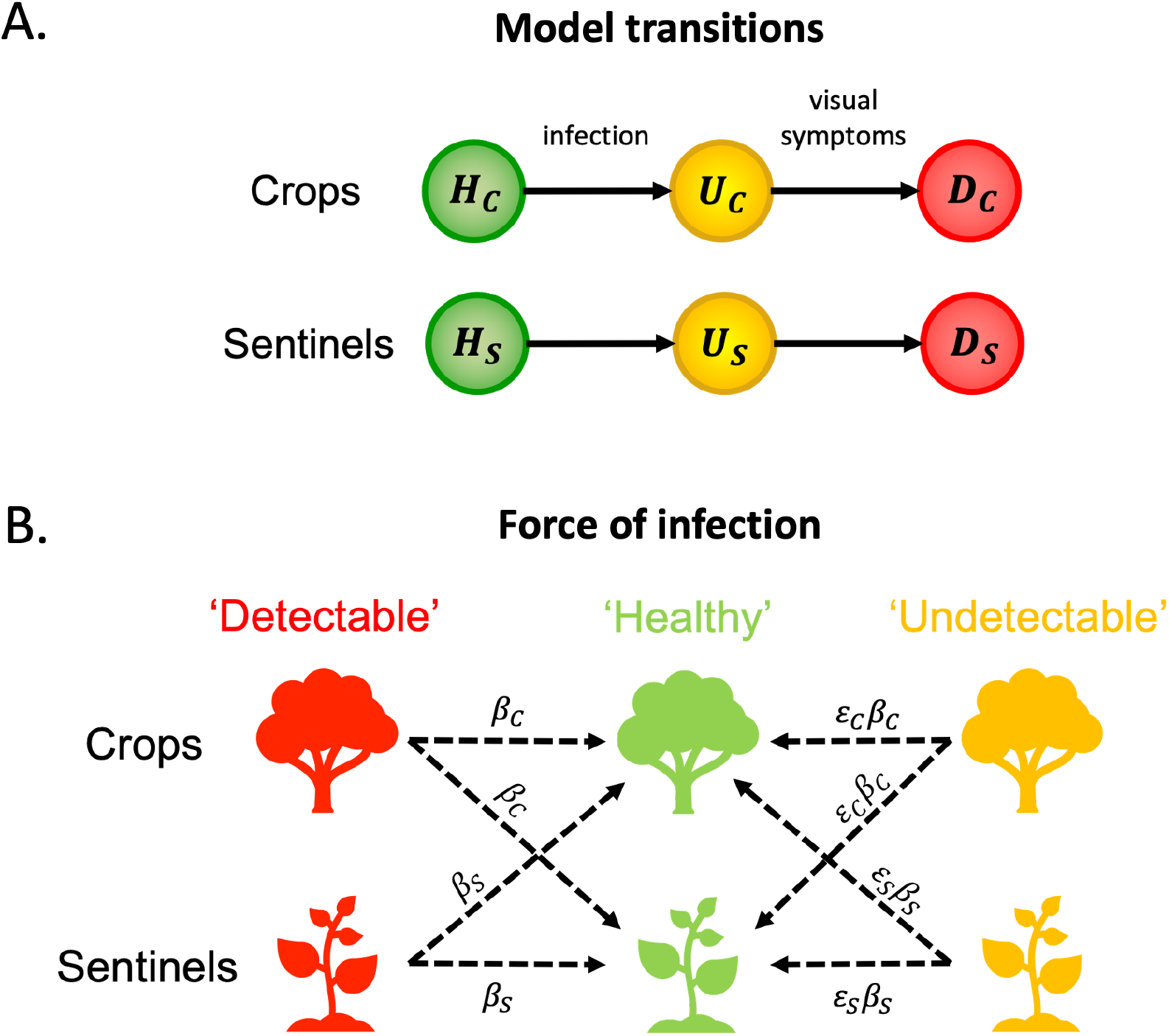
Schematics illustrating the compartments in the model (and how individual plants move between them) and the rates at which infections occur. A. Model transitions. For plants of either type (crop or sentinel), individual hosts begin in the ‘Healthy’ compartment (*H*_*C*_ or *H*_*S*_) before moving to the corresponding ‘Undetectable’ compartment (*U*_*C*_ or *U*_*S*_) upon infection. ‘Undetectable’ plants progress to the appropriate ‘Detectable’ compartment (*D*_*C*_ or *D*_*S*_) once visual symptoms develop. B. Force of infection. The rates at which different infectious hosts generate new infections. A ‘Detectable’ crop plant can infect a ‘Healthy’ crop or sentinel at rate *β*_*C*_ whilst an ‘Undetectable’ crop infects ‘Healthy’ hosts at the scaled rate *∈*_*C*_*β*_*C*_. Similarly, ‘Detectable’ and ‘Undetectable’ sentinels infect ‘Healthy’ plants at rates *β*_*S*_ and *∈*_*S*_*β*_*S*_, respectively.

We denote the total number of crop and sentinel plants in the population by *P*_*C*_ = *H*_*C*_ + *U*_*C*_ + *D*_*C*_ and *P*_*S*_ = *H*_*S*_ + *U*_*S*_ + *D*_*S*_, respectively, with a total population size of *P* = *P*_*C*_ + *P*_*S*_. In each of our model simulations, fixed values of *P, P*_*C*_ and *P*_*S*_ are used, since we consider only the time until first detection and not the subsequent period during which infected plants and other nearby plants may be removed. ‘Undetectable’ and ‘Detectable’ plants may generate new infections in any ‘Healthy’ plant, with crops and sentinels assumed to be equally susceptible. The mode of transmission (insect vectors in the case of *X. fastidiosa*) is captured implicitly through the model parameterisation, rather than modelled explicitly. The parameters *β*_*C*_ and *β*_*S*_ represent the daily per capita rates at which individual infected ‘Detectable’ crop and sentinel plants generate new infections, respectively. We also introduce the scaling parameters *∈*_*C*_ and *∈*_*S*_ to represent the relative infectiousness of ‘Undetectable’ crops and sentinels compared to ‘Detectable’ ones, so that the daily rates at which ‘Undetectable’ crop and sentinel plants generate new infections are *∈*_*C*_*β*_*C*_ and *∈*_*C*_*β*_*C*_ respectively. We assume that ‘Undetectable’ plants are less infectious than ‘Detectable’ ones, so that 0 < *∈*_*C*_, *∈*_*S*_ < 1. The different transmission routes and the corresponding rates at which infections occur are illustrated in Figure 1B. The mean duration of the crop and sentinel ‘Undetectable’ periods are given by the parameters *γ*_*C*_ and *γ*_*S*_ respectively.

The resulting compartmental differential equation model representing pathogen transmission is given by

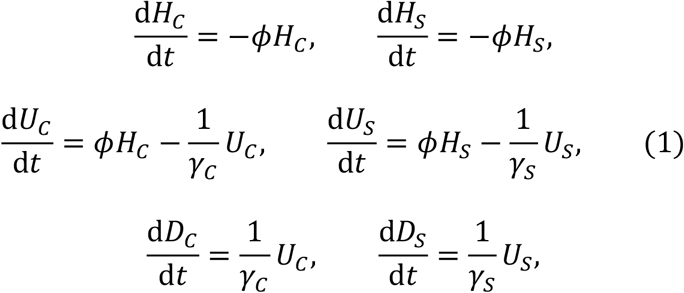

where the force of infection, *ϕ*, acting on each ‘Healthy’ plant is

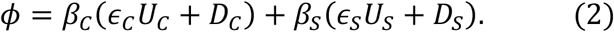

In our analyses, we run simulations of the analogous stochastic model using the direct method version of the Gillespie stochastic simulation algorithm, as described in Section 2.5.

### 2.2 The baseline case – reduced model in the absence of sentinel plants

In the absence of sentinel plants (*P*_*S*_ = 0), the transmission model reduces to

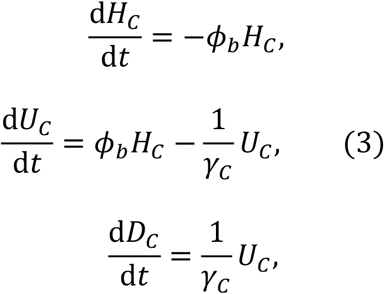

where the force of infection, *ϕ*_*b*_, acting on each ‘Healthy’ plant is

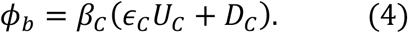

Throughout, this reduced system is what we will refer to as the ‘baseline case’ for a particular choice of model parameterisation or surveillance strategy. It provides a point of reference for the EDP, allowing us to determine whether the EDP is reduced or increased when sentinel plants are introduced. Specifically, the effect of including sentinel plants in a surveillance strategy may then be assessed relative to the baseline case with equivalent parameterisation and sampling effort (i.e. the same total sample size and sample interval). When considering the baseline case, we again run model simulations of the analogous stochastic model using the direct method version of the Gillespie stochastic simulation algorithm.

### 2.3 Sampling and detection

We considered a monitoring programme in which different random samples of *N* hosts are taken from the population every Δ days and inspected for symptoms of disease. We chose random sampling since it outperforms repeated sampling of the same hosts (see Supplementary Text S3 and Supplementary Fig S15), whilst being straightforward to implement computationally. For a given surveillance strategy (choice of *N* and Δ), we fix the number of crop plants and sentinel plants in the sample so that *N* = *N*_*C*_ + *N*_*S*_, where *N*_*C*_ is the crop sample size and *N*_*S*_ is the sentinel sample size (in the baseline case, *N*_*S*_ = 0 and *N*_*C*_ = *N*). To reflect the introduction of the pathogen at a random time relative to the sampling scheme, we begin sampling from our model disease system at a time selected uniformly at random from the interval [0, Δ]. We then sample every Δ days until detection occurs. In our analyses, we assume that ‘Detectable’ plants in a sample are always correctly identified as being infected (no false negatives), ‘Undetectable’ plants are never correctly identified as being infected (no true positives), and ‘Healthy’ plants are always correctly identified as uninfected (no false positives). Therefore, detection occurs at a given time if and only if at least one ‘Detectable’ plant (crop or sentinel) is included in the sample selected at that time.

### 2.4 Model parameterisation

We selected the epidemiological parameters of our model (Table 1) to be consistent with *X. fastidiosa* subsp. *pauca* infection in the crop plant *O. europaea* (European olive) and the sentinel plant *C. roseus* (Madagascan periwinkle) [35, 61]. In the absence of data on transmission in *C. roseus* specifically, in our main analyses we make the conservative assumption (in terms of the derived benefit of using sentinel plants) that the transmission coefficient for ‘Detectable’ sentinels was equal to that for ‘Detectable’ crops (*β*_*C*_ = *β*_*S*_). This is more likely to be an overestimate for *β*_*S*_ than an underestimate; since *C. roseus* is much smaller than *O. europaea*, the rate at which an individual host causes infections is likely to be lower. In choosing this value we therefore provide a lower bound on the utility of sentinel plants. We were also required to assume the relative infectiousness of ‘Undetectable’ sentinels compared to ‘Detectable’ sentinels. The value chosen (*∈*_*S*_ = 0.1) is similarly a conservative choice compared to the equivalent parameter for crop plants (*∈*_*C*_ = 0.015) since it assumes that ‘Undetectable’ sentinels are substantially more infectious than ‘Undetectable’ crops. Again, this choice aims to provide a lower bound on the utility of sentinel plants. Due to the uncertainty in these parameter values, we also performed extensive sensitivity analyses to determine how the choice of parameterisation affected our results (Section 3.4, Supplementary Text S1 and Supplementary Figs S1-14). In each case that we considered, our overall finding - that sentinel hosts can be helpful to reduce the EDP – was unchanged.

**Table 1.** The epidemiological parameters of the model, their meanings, and their baseline values chosen to be consistent with *X. fastidiosa* infection in *O. europaea* (crop) and *C. roseus* (sentinel). Other model parameter values are considered in the Supplementary Material.

| Parameter | Meaning | Value | Justification |
| --- | --- | --- | --- |
| $\beta_c$ | Transmission coefficient for ‘Detectable’ crops | $5 \times 10^{-5}$ | Chosen so that $\beta_c P_C = 0.05$ [61] |
| $\beta_s$ | Transmission coefficient for ‘Detectable’ sentinels | $5 \times 10^{-5}$ | Assumed |
| $\epsilon_c$ | Transmission scaling factor for ‘Undetectable’ crops | 0.015 | [61] |
| $\epsilon_s$ | Transmission scaling factor for ‘Undetectable’ sentinels | 0.1 | Assumed |
| $\gamma_c$ | Duration of crop ‘Undetectable’ period | 452 days | [35] |
| $\gamma_s$ | Duration of sentinel ‘Undetectable’ period | 49 days | [35] |

Throughout, we considered a crop population of *P*_*C*_ = 1000 plants. The size of the sentinel population, *P*_*S*_, was allowed to vary between simulations. We initialised each simulation with a single ‘Undetectable’ infected host, with the probability of that host being a crop or a sentinel plant determined by their respective proportions within the population.

### 2.5 Computational implementation

As noted above, for a given number of sentinels, *P*_*S*_, added to the population, we performed simulations of pathogen spread using the Gillespie stochastic simulation algorithm (direct method) [62], generating stochastic epidemic curves in which the numbers of ‘Undetectable’ and ‘Detectable’ crops and sentinels were tracked over time until the entire population became infected (Fig 2A). Then, given the remaining parameters defining the surveillance strategy (values of *N, N*_*C*_, *N*_*S*_ and Δ), we implemented the corresponding monitoring programme on these simulated epidemics as described in Section 2.3 (Figs 2B,C). For each sampling run and subsequent detection completed on a unique epidemic curve, we recorded the total prevalence of infection in crop plants when the pathogen was discovered (i.e., we recorded the value of *U*_*C*_ + *D*_*C*_ on discovery) (Fig 2C). Repeatedly implementing the same surveillance strategy on many simulated epidemic curves, we obtained a distribution on the discovery prevalence amongst crop plants for that surveillance strategy and computed the EDP as the mean value of that distribution (note that this does not include the prevalence amongst sentinel plants, since we assume that damage to the crop population is the primary concern for commercial growers) (Fig 2D). For any given choice of *N* and Δ, the baseline EDP for that strategy is computed in the same way, setting *P*_*S*_ (and thus also *N*_*S*_) equal to 0.

**Fig 2.**
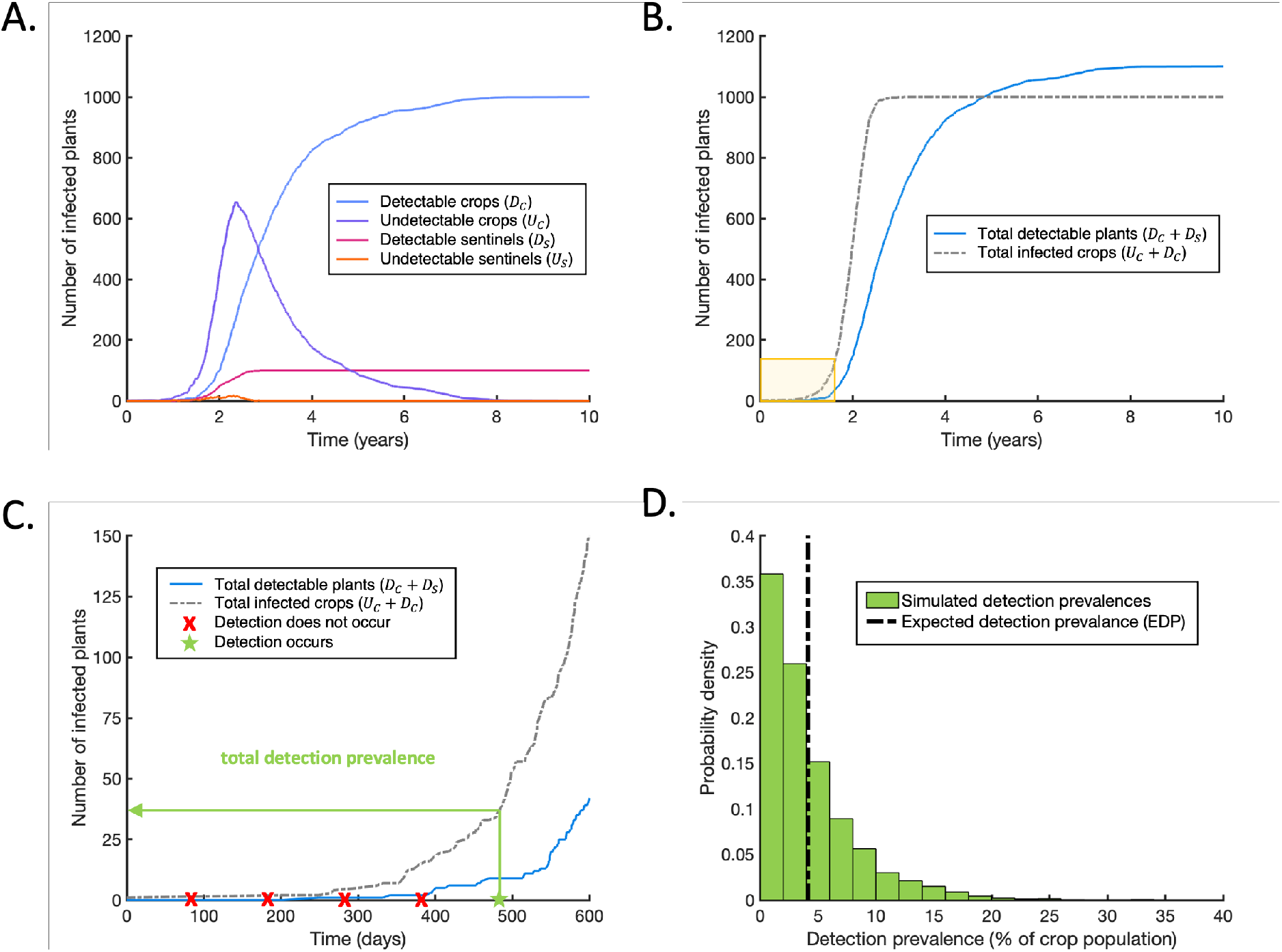
Schematic summarising how the EDP was obtained for an example sampling strategy. A. Stochastic simulations of pathogen spread in the model system were performed using the Gillespie SSA (direct method). In each simulation the number of ‘Undetectable’ and ‘Detectable’ crops and sentinels were tracked over time until the entire population became infected. In the simulation shown, we considered a population of *P*_*C*_ = 1000 crops and *P*_*S*_ = 100 sentinels, beginning with a single ‘Undetectable’ individual. All epidemiological parameters were as specified in Table 1. B. The total number of ‘Detectable’ plants in the population (*D*_*C*_ + *D*_*S*_; blue solid line) and the total number of infected crop plants in the population (*U*_*C*_ + *D*_*C*_; grey dash-dotted line). Yellow box highlights the region that is shown magnified in panel C. C. A magnification of the yellow region highlighted in panel B, showing how the sampling strategy (defined in terms of the parameters *N, N*_*C*_, *N*_*S*_ and Δ) is implemented on an individual epidemic curve. Random samples of size *N* = *N*_*C*_ + *N*_*S*_ were taken from the population every Δ days, with detection occurring as soon as a ‘Detectable’ plant (*D*_*C*_ or *D*_*S*_) was present in the sample. In this example, we used *N*_*C*_ = 50, *N*_*S*_ = 50, Δ = 100 days. The time of the first sample was selected uniformly at random from the interval [0,100]. When detection occurred, the total detection prevalence amongst crop plants (*U*_*C*_ + *D*_*S*_) was recorded. D. The sampling strategy described in C was repeated on 10000 simulated epidemic curves. Due to the inherent randomness of the spread and sampling simulations, different values for the total detection prevalence were obtained each time, leading to a probability distribution for this prevalence (green bars). The EDP was calculated as the mean of this distribution (black dash-dotted line).

Initially, we considered fixed values for the number of sentinels *P*_*S*_ added to the population (Section 3.2). In each case, for a given choice of *N* and Δ we allowed the number of those sentinels included in the sample (*N*_*S*_) to vary, and applied a Bayesian optimisation algorithm ([63, 64]; see also Supplementary Text S2) to identify the choice of *N*_*S*_ for which the EDP was maximally reduced compared to the baseline level. Subsequently, we considered varying the number of sentinels in the population (*P*_*S*_) and in the sample (*N*_*S*_) simultaneously and sought to identify the pair of values 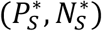 that maximised the reduction in EDP compared to the baseline (Section 3.3). To do so, we adapted the Bayesian optimisation algorithm to allow the objective function to depend on two input variables, and constrained it to ensure that the number of sentinels included in the sample could not exceed the total number of sentinels available in the population (Supplementary Text S2).

Throughout, the EDP for a given surveillance strategy (choice of *P*_*S*_, *N*_*C*_, *N*_*S*_, and Δ) was calculated by performing sampling on 25,000 simulated epidemic curves. All computing code used to implement the above methods was written in MATLAB (compatible with version R2022a), and is available at https://github.com/francescalovellread/sentinel_plants.

## 3 Results

### 3.1 The baseline case – a monitoring programme without sentinel plants

We first considered the effects of implementing a monitoring programme without sentinel plants (the baseline case described in Section 2.2). A schematic of the reduced version of the model system in this case is shown in Fig 3A. In the absence of sentinel plants, the monitoring programme requires selecting a random sample of *N* = *N*_*C*_ plants at regular time intervals Δ and checking for the presence of ‘Detectable’ plants (*D*_*C*_) in the sample (Fig 3B). Doing so, we compute the baseline EDP for sample sizes *N*_*C*_ = 25, 30, 35, …, 200 and sample intervals Δ = 30, 35, 40, …, 150 days (Fig 3C). As expected, lower EDPs are achieved with larger sample sizes *N*_*C*_ (inspecting more plants) and smaller sample intervals Δ (inspecting more frequently). These baseline values provide a point of comparison that we will use to evaluate the relative effects of sentinel-based strategies in subsequent sections.

**Fig 3.**
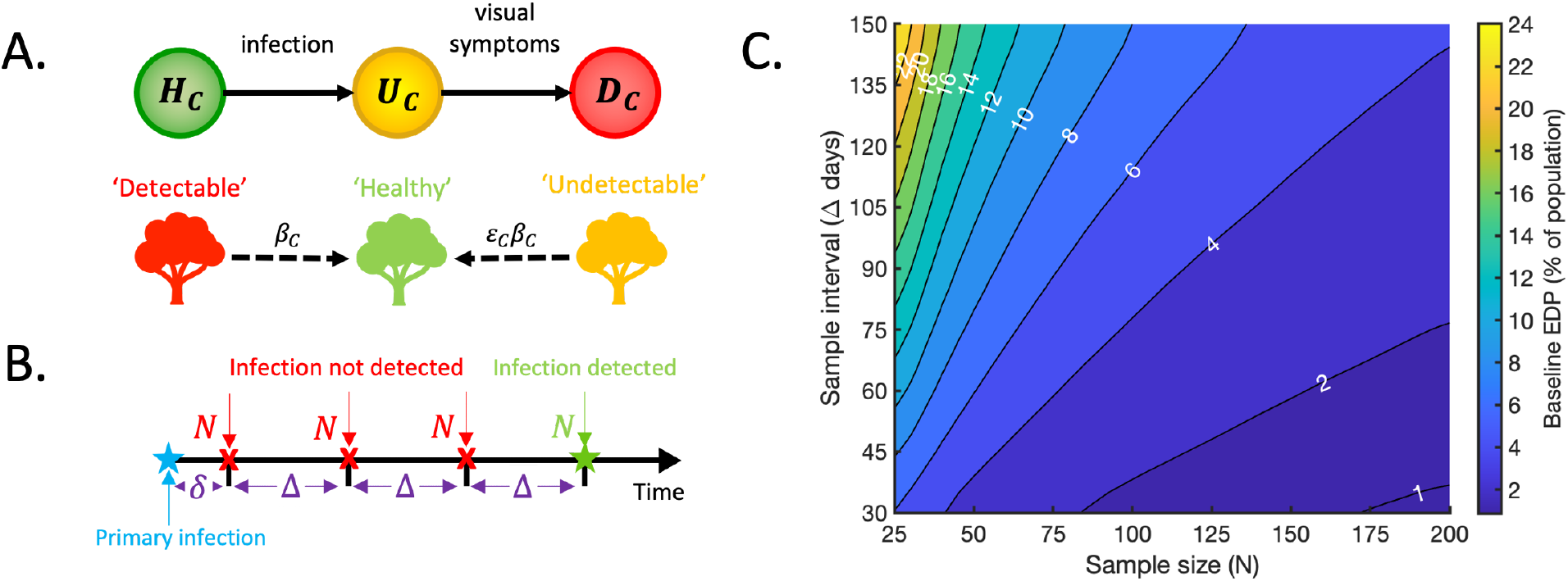
The baseline case – the model in the absence of sentinel plants. A. Schematic illustrating how crop plants progress through the model compartments, and the rates at which transmissions occur. Individual hosts begin in the ‘Healthy’ compartment (*H*_*C*_), move to the ‘Undetectable’ compartment (*U*_*C*_) upon infection and progress to the ‘Detectable’ compartment (*D*_*C*_) once visual symptoms develop. A ‘Detectable’ crop infects ‘Healthy’ crops at per capita rate *β*_*C*_ whilst an ‘Undetectable’ crop generates infections at the scaled per capita rate *∈*_*C*_*β*_*C*_. B. Schematic illustrating the implementation of the monitoring programme. Monitoring begins at a random time δ relative to the time of primary infection, where δ is drawn from a *U*[0, Δ] distribution. Random samples of size *N* are subsequently selected from the population at regular time intervals Δ. Infection is detected at a given time if a ‘Detectable’ plant is contained in the sample selected at that time. C. The baseline EDP, expressed as a percentage of the total crop population size, as the sample size (*N* = *N*_*C*_) and sample interval (Δ) vary.

### 3.2 Introducing sentinel plants – choosing *P*_*S*_ and *N*_*S*_ carefully is critical

We next considered introducing sentinel plants to the population using the full model described in Section 2.1. This raises two important questions.

1. **How many sentinels should we add to the population (***P*_*S*_**)?** Although the relatively fast symptom development of sentinels facilitates the rapid detection of disease, this is only beneficial if the faster discovery time corresponds to a lower EDP. Since adding sentinels will also increase the rate of pathogen transmission, including too many sentinels negates the benefits of fast detection, particularly if (as assumed here) sentinel plants are more infectious than crop plants when ‘Undetectable’.
2. **How many of those sentinels should we include in the sample (***N*_*S*_**)?** Although a natural choice may be to sample preferentially from the available sentinel population (i.e. to include as many sentinels as possible in the sample), this is not necessarily optimal. For example, if the number of sentinels in the population is close to the sample size, this strategy would lead to frequent repeated sampling of the same set of plants, resulting in a reduction in the information gained per sample (see Supplementary Text S3 and Supplementary Fig S15).

In this section, we demonstrate how choosing *P*_*S*_ and *N*_*S*_ carefully is critical to avoid the introduction of sentinel plants having a negative effect and instead achieve the maximum possible reduction in EDP for a given sampling effort.

We began by considering three fixed values for the number of sentinels added: *P*_*S*_ = 50, *P*_*S*_ = 100 and *P*_*S*_ = 200. We allowed the total sample size *N* to take values *N* = 25, 50, 75, …, 200, and considered five values for the sample interval: Δ = 30, 60, 90, 120 and 150 days. In each case, we allowed the number of sentinels included in the sample (*N*_*S*_) to be chosen in the range [0, min(*P*_*S*_, *N*)]. This choice of upper limit ensures that the number of sentinels sampled does not exceed the total sample size (*N*) or the total number of sentinels available (*P*_*S*_).

For each combination of (*P*_*S*_, *N*, Δ), we ran the Bayesian optimisation algorithm (see Section 2.5 and Supplementary Text S2) to identify the choice of *N*_*S*_ corresponding to the greatest reduction in EDP compared to the baseline value for that (*N*, Δ) pair. We denoted this optimal choice of *N*_*S*_ by 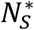. For example, in the case *P*_*S*_ = 50, *N* = 50, and Δ = 30 days, the optimisation indicated that the maximum reduction in EDP was achieved when 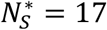 sentinels were included in each sampling round (out of a total possible maximum of 50) (Fig 4A). This choice of sampling strategy (indicated by the green circle) led to a 16% reduction in the EDP compared to the baseline value. When *N*_*S*_ was instead chosen to take another of the values considered, smaller reductions (or even increases) in the EDP were achieved. In the extreme case in which none of the available sentinels were sampled, an 11% increase in the EDP was observed (y-intercept of Fig 4A), demonstrating the drawback of adding sentinel plants to the population (an increase in transmission rate) if they are not sampled appropriately. The optimal number of sentinels 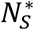 to include in the sample across the range of sample sizes (*N*) and sample intervals (Δ) is shown for *P*_*S*_ = 50, 100 and 200 in Figs 4B,C,D respectively, with the corresponding reductions in the EDP compared to the baseline shown in Figs 5A,B,C.

**Fig 4.**
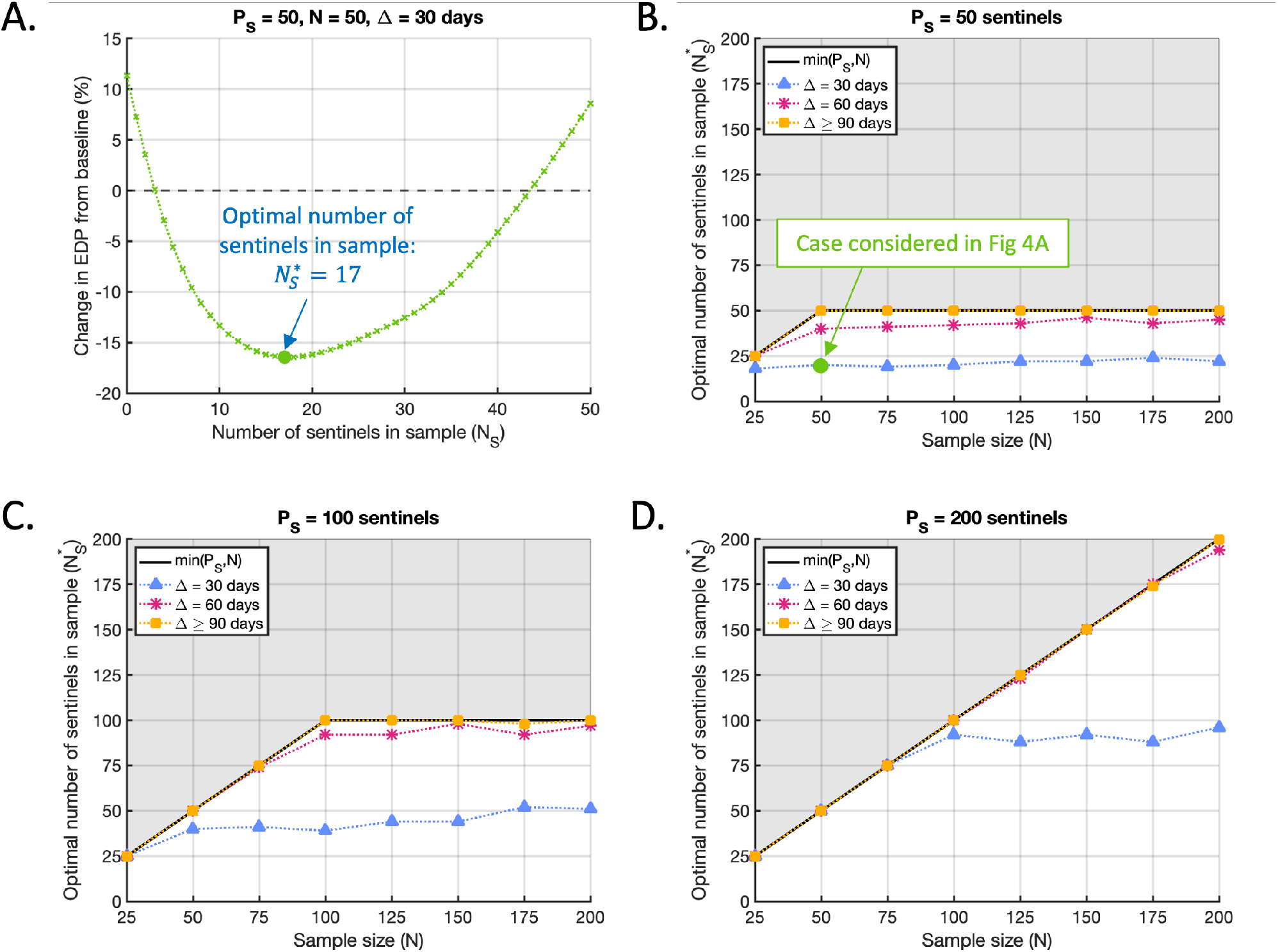
The optimal number of sentinel plants to include in the sample depends on the sample size, sample interval and the total number of sentinels in the population. A. The effect of varying the number of sentinels included in the sample (*N*_*S*_) on the percentage change in EDP compared to the baseline level, in the case *P*_*S*_ = 50, *N* = 50, Δ = 30 days. The number of sentinels in the sample for which the reduction in EDP is maximised 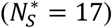 is indicated by the green circle. Black dashed line marks the baseline EDP. B. The optimal number of sentinels 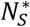 to include in the sample when *P*_*S*_ = 50, as the sample size (*N*) and sample interval (Δ) vary. Solid black line marks the maximum possible number of sentinels that could be sampled at any time (min*P*_*S*_, *N*)). Grey shading marks the unfeasible region in which *N*_*S*_ exceeds this maximum. Green circle marks the case considered in A (*P*_*S*_ = 50, *N* = 50, Δ = 30 days). C. The analogous figure to B, but with *P*_*S*_ = 100 sentinels added to the population. D. The analogous figure to B, but with *P*_*S*_ = 200 sentinels added to the population.

**Fig 5.**
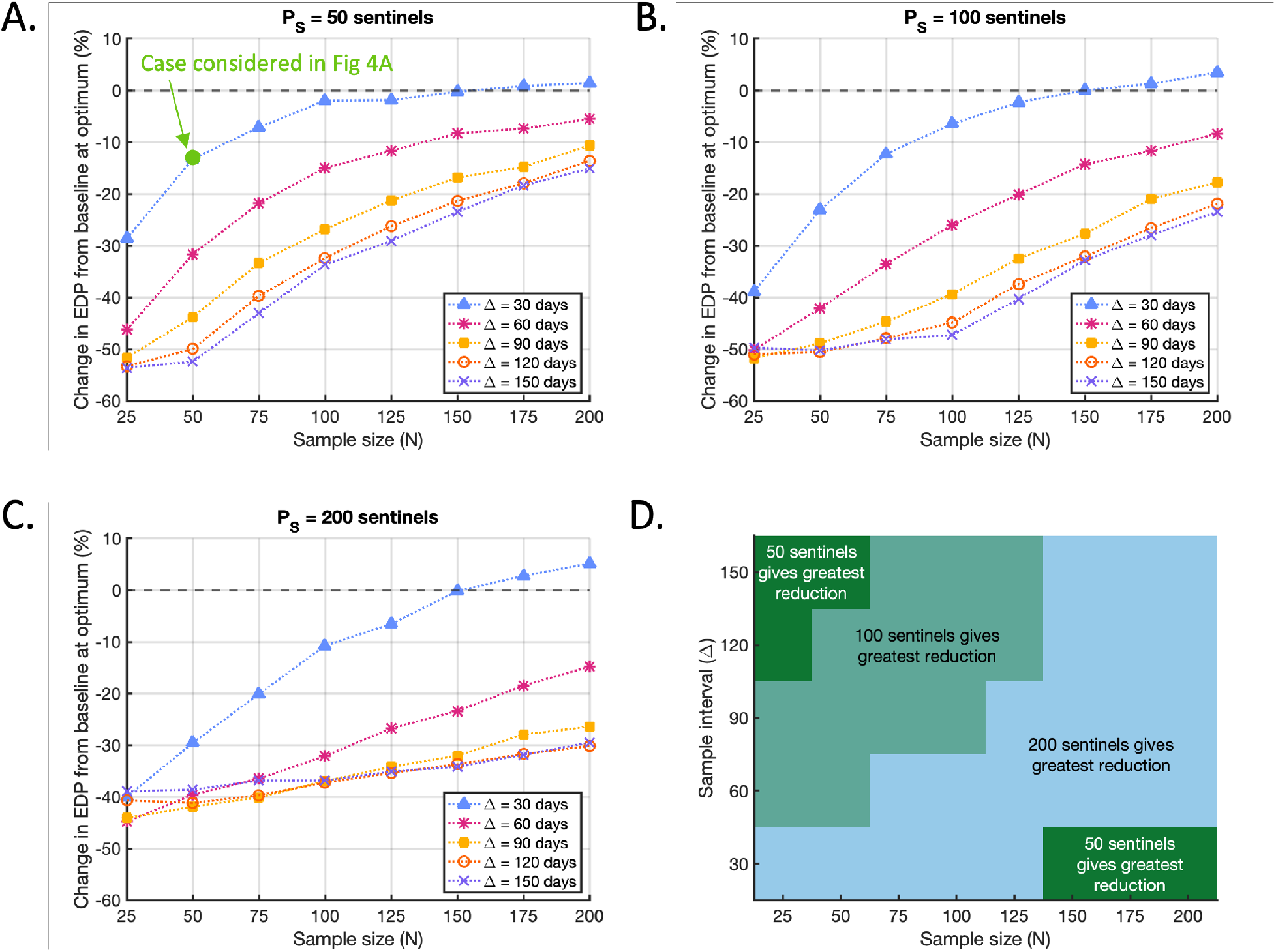
Optimal reductions in EDP compared to the baseline level. A. The best achievable percentage changes in the EDP compared to the baseline level for each (*N*, Δ) pair when *P*_*S*_ = 50, corresponding to the optimal strategies identified in Fig 4B. Green circle marks the case considered in Fig 4A (*P*_*S*_ = 50, *N* = 50, Δ = 30 days). Note that the baseline level depends on *N* and Δ (Fig 3C, Supplementary Fig S16A), so the relative changes in EDP shown here are not a measure of the resultant EDP. The resultant EDP decreases with sampling effort (Supplementary Figs S16B,C,D). B. The analogous figure to A, but with *P*_*S*_ = 100 sentinels added to the population and results corresponding to the strategies identified in Fig 4C. C. The analogous figure to A, but with *P*_*S*_ = 200 sentinels added to the population and results corresponding to the strategies identified in Fig 4D. D. Combinations of the sample size *N* and sample interval Δ for which adding *P*_*S*_ = 50 (dark green), *P*_*S*_ = 100 (light green) or *P*_*S*_ = 200 (blue) sentinels to the population led to the greatest reduction in the EDP compared to the baseline level (of the three values of *P*_*S*_ considered).

The optimal number of sentinel plants to include in the sample depended strongly on the sample interval and on the relationship between the sample size and the total number of sentinels available (Figs 4B,C,D). When Δ = 90 days, 120 days or 150 days, the optimal strategy in every case we considered was to sample the maximum possible number of sentinels (i.e. 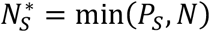). Since this result is identical for all three of those cases, they are represented by the single yellow line in Figs 4B,C,D. However, for the shorter sample intervals of Δ = 30 and 60 days (blue and pink lines respectively), the optimal monitoring strategy involved sampling a combination of sentinel plants and crop plants. In other words, in those scenarios it was preferable to sample fewer than the maximum allowable number of sentinels (i.e. 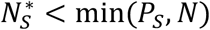) for a range of choices of *P*_*S*_ and *N*, particularly when the total number of sentinels in the population was not substantially larger than the sample size.

These results may be explained by noting that, if the total number of sentinels available to sample from (*P*_*S*_) is not substantially larger than the sample size (*N*), then sampling the maximum allowable number of sentinels (min*P*_*S*_, *N*)) results in many or all of the same plants being repeatedly selected in every sampling round. If the sample interval is short, this leads to the frequent re-inspection of plants whose disease-free status was already established in the recent past, limiting the information gained per sampling round. However, this effect diminishes as the sample interval increases, because the disease status of plants inspected in the previous sample becomes less informative of their state at the next sample time. Thus, when the sample interval is large, sampling the maximum possible number of sentinels is the optimal strategy 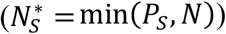 regardless of the sample size (*N*) or the total number of sentinels available (*P*_*S*_). These results confirm the need to consider the division of the sample between crops and sentinels as a variable quantity that should be chosen carefully based on the precise conditions under which surveillance is taking place. If the number of sentinels included in the sample is suboptimal, smaller reductions in the EDP will be achieved, and sentinel plants may even have a detrimental effect (Fig 4A).

For almost all values of *P*_*S*_, *N* and Δ that we considered, when the number of sentinels included in the sample was optimised (as indicated in Fig 4), a reduction in the EDP compared to the baseline value was achieved (Figs 5A,B,C). Exceptions to this occurred when the sample interval was Δ = 30 days and the sample size was *N ≥* 150: in those cases we observed a slight increase in the EDP compared to the baseline (Figs 5A,B,C). This can be attributed to the fact that, in this initial analysis, we considered only a limited number of values of *P*_*S*_ (specifically, *P*_*S*_ = 50, 100 or 200). If instead *P*_*S*_ can be chosen to take any value, the baseline EDP would never be exceeded at the optimised value of *P*_*S*_, as at worst the baseline EDP can always be achieved by setting *P*_*S*_ = 0. These results demonstrate that, in addition to the proportion of sentinels included in the sample, the total number of sentinels added to the population must also be carefully selected: if a suboptimal value of *P*_*S*_ is chosen, sentinel plants can be detrimental rather than beneficial.

As expected, the resultant EDP following the implementation of the optimal sentinel strategy decreased with greater sampling effort: taking larger samples and/or sampling more frequently always led to a lower EDP (Supplementary Fig S16). However, larger percentage reductions in the EDP relative to the baseline level were mostly achieved when the sampling effort was low (i.e. when the sample size was small and/or the sample interval was large) (Figs 5A,B,C). This is because, when the sampling effort was low, the baseline EDP was much higher to begin with (Fig 3C). In those cases, the potential for the use of sentinel plants to lead to a large relative improvement in the EDP was greater than when the sampling effort was high and the baseline EDP was already low.

As well as affecting the magnitude of the reduction in EDP compared to the baseline, the choice of sample size and sample interval also affected the total number of sentinels for which the greatest reduction was achieved (Fig 5D). For example, when the sample size was *N* = 25 and the sample interval was Δ = 150 days, choosing *P*_*S*_ = 50 led to the greatest reduction in EDP of the three values considered (54%, compared to a 50% reduction when *P*_*S*_ = 100 and a 39% reduction when *P*_*S*_ = 200). However, for *N* = 100 and Δ = 30 days, choosing *P*_*S*_ = 200 gave the greatest reduction in EDP (11%, compared to 2% and 6% when *P*_*S*_ = 50 and 100, respectively). Overall, introducing fewer sentinels was preferable when the sampling effort was either low or very high, with larger numbers preferable for intermediate sampling efforts (Fig 5D).

This variation in the optimal number of sentinels for different values of (*N*, Δ) reflects the crucial trade-off between the benefits and drawbacks of sentinel plants. Although adding sentinels to the population helps to facilitate early detection, it also leads to an increased rate of pathogen transmission (particularly if sentinels are more infectious than crop plants when ‘Undetectable’, as assumed here). Therefore, including more sentinel plants is only beneficial if the advantage gained from sampling them outweighs the impact of increased transmission.

For small sample sizes *N*, the capacity to exploit large numbers of sentinel plants is limited. Although increasing the number of sentinels is beneficial up to a point, since it allows for sampling without frequently inspecting the same sentinel plants, there is a threshold number of sentinels to introduce beyond which there will be no further improvement in detection to counterbalance the concurrent increase in overall transmission. The benefit of increasing *P*_*S*_ is more limited when the sample interval Δ is large, since in that scenario a past negative sample is less likely to indicate that the current sample will be negative. Thus, the same sentinels may be resampled without a substantial correlation between successive samples. Smaller numbers of sentinels are therefore preferable when the sampling effort is low (Fig 5D). At the opposite extreme, when sampling is very intensive (large sample size *N* and small sample interval Δ) then the baseline EDP is low (Fig 3C) and the potential for sentinel plants to reduce it is limited. In such a case, including a very large number of sentinels in the population is also not optimal, since this limited reduction is outweighed by the consequent higher rate of transmission. Therefore, smaller numbers of sentinels are also preferable when the sampling effort is very high (Fig 5D). However, for intermediate sampling efforts, larger numbers of sentinels perform better, since the capacity to exploit them and the scope to reduce the EDP compared to the baseline are less restricted. These results emphasise that judicious selection of the total number of sentinel plants added to the population is required to ensure that the benefits of including them are sufficient to offset their drawbacks in terms of increasing transmission. This emphasises the need for an epidemiological modelling framework as provided here to guide the number of sentinel plants to introduce, and we explore how the number of sentinel plants can be optimised in the next section.

### 3.3 Optimising the number of sentinel plants included in the population

We next considered optimising the total number of sentinel plants added to the population (*P*_*S*_) and the number of sentinels included in the sample (*N*_*S*_) simultaneously, in order to achieve the maximal reduction in the EDP. We allowed the total sample size *N* to take values *N* = 25, 30, 35, …, 200, and the sample interval to take values Δ = 30, 35, 40, …, 150 days. In every case, we allowed the total number of sentinels added to the population (*P*_*S*_) to vary in the range [0,350], and the number of sentinels included in the sample (*N*_*S*_) to vary in the range [0,min*P*_*S*_, *N*)]. The upper bound of 350 on *P*_*S*_ was selected following trial simulations that indicated this would be sufficient to identify the optimal value of *P*_*S*_ across the (*N*, Δ) range considered. For each (*N*, Δ) pair, we applied the Bayesian optimisation algorithm on *P*_*S*_ and *N*_*S*_ (constrained such that *N*_*S*_ ≤ *P*_*S*_; see Supplementary Text S2) to find the pair of values 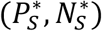 that reduced the EDP most compared to the baseline value.

Including sentinel plants in the population was beneficial across the range of sampling strategies considered, with the optimal number of sentinels added to the population 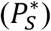 greater than zero for all values of (*N*, Δ) (Fig 6A). This shows that deploying sentinel plants has the potential to reduce the EDP. However, the optimal number of sentinels to use varied substantially with the sample size (*N*) and sample interval (Δ). Consistent with Fig 5D, 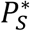 was low when the sampling effort was small (region 1 in Fig 6A), with the optimal number of sentinels increasing for larger sample sizes and smaller sample intervals. This is again due to the benefit of avoiding repeated sampling of the same plants when the sampling effort is high, as repeatedly sampling the same plants leads to correlation between samples and therefore reduces the amount that is learnt from each sample. By including more sentinels, the chance of repeatedly sampling the same sentinel plants is reduced. As in Fig 5D, 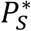 dropped again when the sampling effort was very high (region 2 in Fig 6A). In that region, the baseline EDP was very low (Fig 3C), and the scope for reducing it insufficient to offset the increase in transmission rate caused by adding large numbers of sentinel plants into the population. When the sample interval (Δ) was large, the optimal number of sentinel plants to include in the population was equal to the sample size (*N*) (region 3 in Fig 6A in which contour lines are vertical). In that region, the sample interval was long enough to allow for repeated sampling of the same plants, eliminating the need for *P*_*S*_ to exceed the sample size.

**Fig 6.**
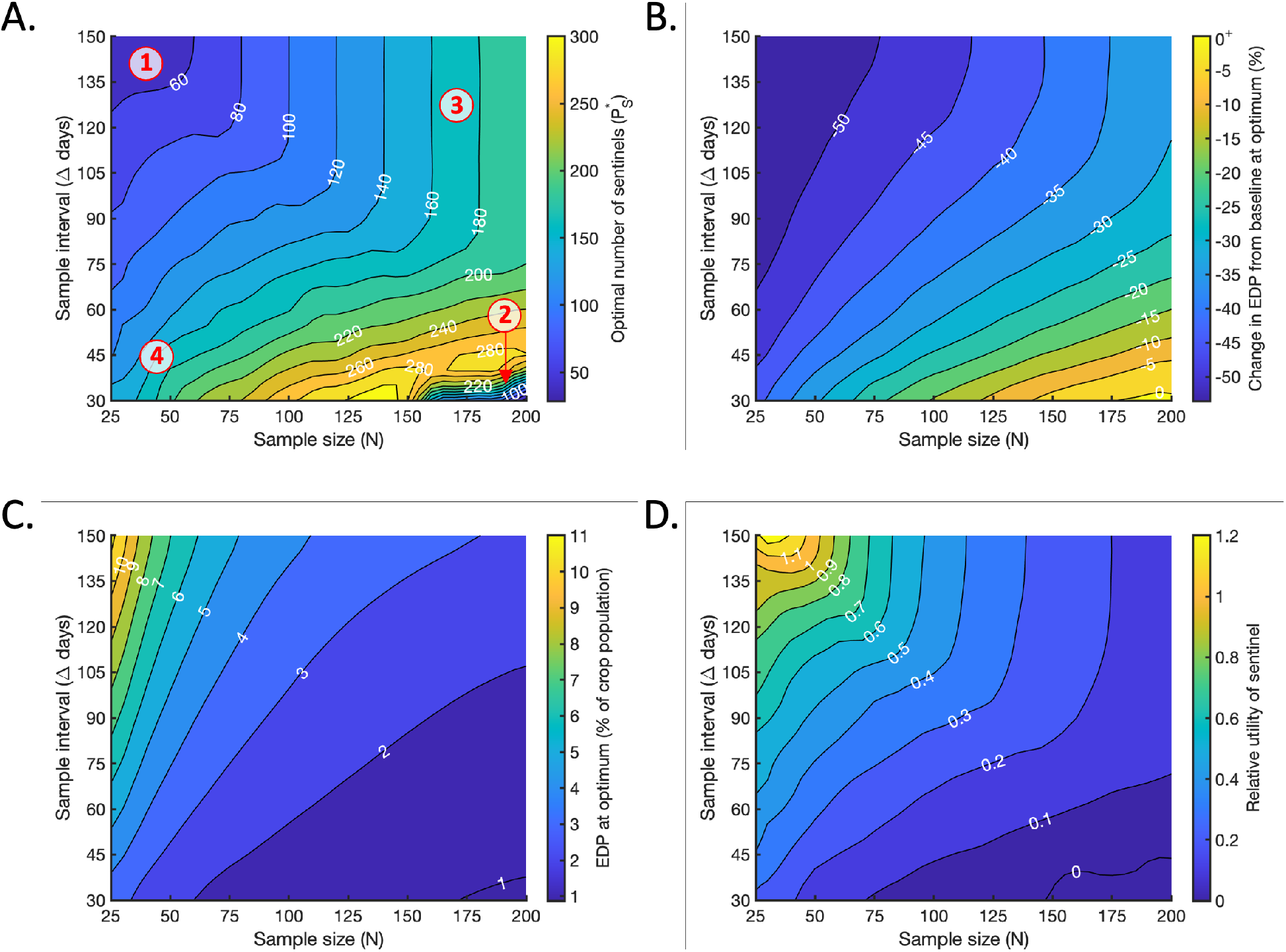
Optimising the number of sentinels to include in the population. A. The optimal number of sentinel plants to include in the population, 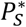, for which the maximal reduction in the EDP compared to the baseline level is achieved (if *N*_*S*_ is also chosen optimally). B. The percentage change in the EDP compared to the baseline value at the optimum, achieved when 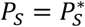 and 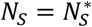. The resultant value of the EDP at the optimum, expressed as a percentage of the total crop population. D. The relative utility of a single sentinel plant (the percentage reduction in EDP per sentinel) at the optimum.

For almost all of the (*N*, Δ) values considered, the optimal number of sentinels to include in the sample 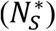 was the maximum possible (i.e. 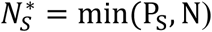) (Supplementary Fig S17). As observed in Section 3.2, preferential sampling of sentinel plants was always optimal when the sample interval (Δ) was large, since the same plants could be sampled repeatedly while still gaining new information about pathogen transmission each time. Sampling the maximum possible number of sentinels was also optimal when the sample interval and sample size were both small. In that region, the optimal total number of sentinels in the population was substantially larger than the sample size (region 4 in Fig 6A), meaning that preferential sampling of sentinels did not result in repeated sampling of the same plants. 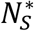 only fell below min(*P*_*S*_, *N*) when the sampling effort was very high (large *N* and small Δ). In that region, the total number of available sentinels dropped below the sample size (region 2 in Fig 6A) and the issue of repeated sampling again became relevant. However, since the sample size substantially exceeded the number of sentinels in that region, repeated sampling had a relatively small effect and the consequent reduction in the optimal proportion of sentinels to sample was not large (Supplementary Fig S17).

When *P*_*S*_ and *N*_*S*_ were chosen optimally, reductions in the EDP compared to the baseline value were achieved for almost all (*N*, Δ) values considered (Fig 6B). Of course, since the baseline itself (Fig 3C) can theoretically always be achieved by choosing *P*_*S*_ = 0, we would not expect the optimal resultant EDP to exceed the baseline substantially for any choice of *N* and Δ. However, in cases where the optimal sentinel strategy has little effect on the EDP, small increases in the EDP compared to the baseline may still occur due to the stochasticity of our simulations. This was observed for some very large values of *N* and very small values of Δ (Fig 6B).

As in all previous cases, the resultant EDP decreased with greater sampling effort: increasing the sample size (*N*) and/or decreasing the sample interval (Δ) always led to a smaller EDP (Fig 6C). However, consistent with Figs 5A,B,C, although the resultant EDP was smallest when the sampling effort was high, larger percentage reductions in the EDP compared to the baseline level were achieved when the sampling effort was low (small *N* and/or large Δ) (Fig 6B). Smaller percentage reductions compared to the baseline were achieved for greater sampling efforts. As noted in Section 3.2, this can be explained by observing that the baseline EDP was much greater for low sampling efforts than high, providing greater scope for relative improvement (Fig 3C). As a result, the relative impact of a single sentinel plant on the reduction in EDP compared to the baseline varied substantially with sampling effort (Fig 6D). The relative utility of a sentinel, computed as the percentage reduction in EDP from the baseline at the optimum (shown in Fig 6B) divided by the optimal number of sentinels to include in the population (shown in Fig 6A), was greatest when the sampling effort was very small, and large reductions in the EDP were achieved using a small number of sentinel plants (Figs 6A,B,C). The relative utility decreased for larger sampling efforts, when more sentinels were included in the population but the reduction in EDP compared to the baseline was less. In practical terms, this suggests that the ‘value for money’ of a single sentinel plant decreases for larger sampling efforts. However, exploring this implication fully would require setting these results in a wider economic context (see Discussion), which is beyond the scope of this study.

### 3.4 Robustness of the results to the parameter values used

As far as possible, the epidemiological parameters used in our analyses were chosen based on literature estimates for *X. fastidiosa* infection in *O. europaea* and *C. roseus* (Table 1). However, reported estimates were not available for all parameters. In particular, we were required to assume values for the transmission coefficient for ‘Detectable’ sentinels (*β*_*S*_) and the transmission scaling factor for ‘Undetectable’ sentinels (*∈*_*S*_) (see Section 2.4). Therefore, we also conducted supplementary analyses to determine how variation in the model parameters affected our results. As well as varying *β*_*S*_ and *∈*_*C*_, we considered variation in the transmission scaling factor for ‘Undetectable’ crops (*∈*_*C*_), the durations of the crop and sentinel ‘Undetectable’ periods (*γ*_*C*_ And *γ*_*S*_), the crop population size (*P*_*C*_) and the initial number of ‘Undetectable’ infected individuals (*U*_0_). For each of these, we generated plots analogous to Fig 6 for two different values of the relevant parameter (Supplementary Text S1 and Figs S1-14).

Although the optimal total number of sentinel plants to include in the population (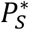; panel A in each of Supplementary Figs S1-14) exhibited some variation with the changes in parameter values, the overall results remained qualitatively similar in most respects. In half of the cases that we considered, the main qualitative change was that the drop in 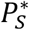 observed in region 2 of Fig 6A did not occur (Supplementary Figs S1,3,4,9,12,13,14). In general, this arose in cases in which sentinel plants became relatively more beneficial due to the parameter change – for example, when the duration of the sentinel ‘Undetectable’ period (*γ*_*S*_) or the relative infectiousness of ‘Undetectable’ sentinels (*∈*_*S*_) was reduced. In some other cases, when the parameter change led to sentinel plants becoming less beneficial, such as increasing the transmission coefficient for ‘Detectable’ sentinels (*β*_*S*_) or the relative infectiousness of ‘Undetectable’ sentinels (*∈*_*S*_), then smaller numbers of sentinels were preferable (Supplementary Figs S2,6,8,10). The optimal number of sentinels was also predictably reduced when the crop population size was halved (*P*_*C*_ = 500; Supplementary Fig S11), confirming that sentinel-based strategies must be assessed in context and tailored to the specific crop population being considered.

As expected, larger percentage reductions in the EDP compared to the baseline were observed for parameter changes that increased transmission amongst crop plants or that made sentinel plants relatively more beneficial (panel B in Supplementary Figs S1,3,4,5,9,13,14). This included increasing the relative infectiousness of ‘Undetectable’ crops, increasing the initial number of infected hosts, and decreasing the infectiousness of sentinels or the duration of their ‘Undetectable’ period. Similarly, for parameter changes that made sentinel plants relatively less beneficial, such as increasing the duration of their ‘Undetectable’ period or their relative infectiousness, smaller reductions in the EDP were observed (Supplementary Figs S2,6,10).

The resultant EDP at the optimum (panel C in each figure) and the relative utility of a single sentinel plant (panel D in each figure) remained qualitatively similar with variation in each parameter. In every case, increasing the sampling effort (increasing the sample size *N* and/or reducing the sample interval Δ) led to a smaller resultant EDP and a reduction in the relative utility per sentinel. Note that for some parameter changes, the optimal number of sentinels fell to 0 when the sampling effort was high, meaning that the sentinel utility was undefined: in those instances, we set the utility to 0 to reflect the fact that sentinel plants had no beneficial effect. The resultant EDP remained quantitatively similar for almost all parameter changes, with the only substantial change occurring when the transmission scaling factor for ‘Undetectable’ crops was increased from *∈*_*C*_ = 0.015 to *∈*_*C*_ = 0.25 (Supplementary Fig S4C).

Despite some variation in the optimal total number of sentinels to include in the population and the magnitude of the corresponding changes in the EDP, in every case that we considered our main conclusions were unchanged. Our results indicate that sentinel plants are often effective at reducing the EDP compared to the baseline level for a range of sampling efforts (Fig 6B and panel B in Supplementary Figs S1-14). They also highlight that the optimal choice of sentinel-based strategy varies according to the sampling resources available (Fig 6A and panel A in Supplementary Figs S1-14). Although higher sampling efforts always led to a lower resultant EDP (Fig 6C and panel C in Supplementary Figs S1-14), in practice the maximum sample size and minimum sample interval a grower can choose will be limited by economic and/or logistic constraints, so that very large sample sizes and small sample intervals may not be feasible. Our model provides a framework for growers to determine how sentinel plants could be used to minimise the resultant EDP subject to their own sampling constraints.

## 4. Discussion

An important challenge in plant disease management is to detect invading pathogens before they become widespread in the host population. In this article, we have shown how early detection of invasive pathogens can be aided by the introduction of sentinel plants, which are alternative hosts that display symptoms of infection more quickly than the main host species. We have explored the trade-off between faster detection of the pathogen using sentinel plants and the potential for sentinel plants to increase pathogen transmission, which is likely if the onset of symptoms is associated with high infectivity. Although we used *X. fastidiosa* infection in *O. europaea* (crop) and *C. roseus* (sentinel) as a case study, we provide a general and extensible epidemiological modelling framework for optimising the deployment of sentinel plants in a broad range of host-pathogen systems.

Our modelling approach involves a compartmental transmission model including two host species (crops and sentinels), in which infected plants of either type are first asymptomatic (‘Undetectable’) before developing visible symptoms. We investigated the impacts of sentinel-based surveillance strategies by implementing a range of different monitoring programmes on simulated epidemic curves. For each monitoring programme considered, we measured the reduction in the EDP compared to a non-sentinel based strategy of equivalent sampling effort (the baseline level; Fig 3C). We used our model to assess the effectiveness of sentinel-based strategies over a range of sample sizes *N* and sample intervals Δ, allowing both the number of sentinels added to the population and the number included in the sample to vary. Overall, our results indicated that sentinel plants have the potential to facilitate substantial reductions in the EDP compared to the baseline level for a wide range of sampling efforts (Figs 5A,B,C, 6B).

The practical benefits of a reduced EDP are multiple. First, discovering the pathogen at a low prevalence increases the chance that local eradication is feasible. Second, a lower prevalence of infection on pathogen discovery minimises the number of plants that must be removed, with benefits including lowering the total value of damaged crops, a lower reduction in crop yield and a reduced logistical cost of plant removal. Third, fast discovery of an invading pathogen limits the risk of dispersal to other locations [14]. Therefore, incorporating sentinel plants into surveillance programs may be beneficial for increasing the chance of success of local control, reducing its cost, and disease containment on a broader scale.

In addition to the overall potential of sentinel plants to facilitate early detection, our results highlight the importance of carefully selecting the number of sentinels planted among the crop population, and the number of sentinels to sample, to ensure that the greatest benefits are achieved. We showed that the optimal number of sentinel plants to include in the population varies with the sample size *N* and sample interval Δ, with smaller numbers of sentinels performing better when the sampling effort was low or very high, and larger numbers of sentinels performing better for intermediate sampling efforts (Figs 5D, 6A). We also showed that it is not always best to focus surveillance resources on sentinel plants alone, as might have been expected when surveillance resources are in short supply. Instead, sampling a combination of both sentinel plants and the main crop can be the optimal strategy, depending on both the sample interval and the size of the sample relative to the total number of sentinels in the population (Fig 4). In particular, sampling as many sentinels as possible is not the best strategy if this results in repeated sampling of the same set of plants, since sampling effort is wasted by re-inspecting plants whose disease-free status has been established in the recent past. Our findings demonstrate the need to consider the number of sentinel plants to introduce, as well as the number of sentinels to sample. The deployment of sentinel plants can be substantially less beneficial, or even detrimental, if either of these quantities are chosen without carefully assessing the impact of the monitoring programme in advance (Figs 4A, 5A,B,C).

The magnitude of the reduction in EDP that was achieved also varied with the choice of sample size and sample interval (Fig 6B). The greatest reductions compared to the baseline were observed for the smallest sampling efforts (small *N* and/or large Δ), when the baseline EDP was high and the scope for improvement large (Fig 3C). In contrast, much smaller reductions relative to the baseline were achieved when the sampling effort was high (Fig 6B), even when greater numbers of sentinels were required to achieve the optimal reductions. Therefore, the relative utility of a sentinel plant (which we defined here to be the percentage reduction in the EDP compared to the baseline level per sentinel in the population) dropped substantially with increased sampling effort (Fig 6D). Although here we do not attempt to quantify the economic impacts of using sentinel plants as a tool for early detection monitoring, this result suggests that careful consideration is needed to evaluate scenarios in which the use of sentinel plants provides good value for money. In this study, our focus was investigating the degree to which including sentinel plants in a surveillance programme could reduce the EDP, thereby increasing the feasibility of pathogen containment and reducing the cost of post detection control. However, although our analyses suggest that sentinel plants may be effective for this purpose, there is clear motivation to consider these results within a wider economic context for specific pathogens and for different crop and sentinel hosts. In practice, an important consideration is the cost of implementing the surveillance programme, which must be compared against the benefits of fast detection of the invading pathogen. Introducing and maintaining sentinel plants will incur additional expense, and may not be cost effective when the utility per sentinel is low. Further research is therefore needed to explore the trade-off between the economic costs of surveillance and control, accounting for the cost of sampling as well as the cost of removing infected plants and the resulting loss in plant value. The specific objective of the control strategy also needs to be carefully considered. For example, if reducing the risk of the pathogen being exported to a new location prior to detection is a particular focus, the decision maker may wish to consider the ‘global’ epidemic cost associated with pathogen exportation in addition to the ‘local’ cost incurred in the controlled region [20, 26], or to seek surveillance strategies that minimise the detection prevalence in the sentinel population as well as the crop population (for a preliminary analysis, see Supplementary Text S4 and Fig S18).

Due to uncertainty in the values of the epidemiological parameters used in our model, we also conducted supplementary analyses to assess how our results varied when these parameters were altered (Supplementary Text S1 and Figs S1-14). Although we observed some quantitative variation in the optimal total number of sentinels to include in the population and the corresponding reduction in the EDP compared to the baseline, our main conclusion was unchanged: sentinel plants can be beneficial for reducing the EDP across a broad range of sampling efforts and model parameters. However, the optimal sentinel strategy varies with the sampling resources deployed, as well as with the underpinning epidemiology of the system under consideration. Therefore, different sentinel-based strategies should be carefully assessed to ensure that the maximum benefits are achieved for a given sampling effort.

Although here we considered *X. fastidiosa* infection in *O. europaea* and *C. roseus* as a case study, our model provides a general framework that can be used to assess sentinel-based surveillance strategies in other pathosystems. For example, monitoring programs for citrus greening disease (a bacterial infection of citrus with causal agents *Candidatus* Liberibacter spp.) are similarly hindered by a long period of asymptomatic infection in mature trees [37, 38]. However, younger trees develop symptoms more quickly [37, 42], and our model could be adapted to consider their use as sentinel plants in this case. In addition to an appropriate parameterisation, some other model adjustments may be necessary when considering an alternative pathosystem. For example, when considering citrus greening disease in a commercial plantation, young sentinel trees may have to replace older trees rather than being interspersed amongst them due to their size and the harvesting equipment used. In that scenario, including sentinel plants could have a detrimental effect on the crop yield, and this must be considered alongside the potential benefits of early detection.

Our aim here was to use a simple model to gain broad insights into the potential of sentinel plants to facilitate early detection. However, our model could be extended to incorporate additional realism in several ways. For example, we could extend the transmission model considered here to account for the spatial structure of the population, allowing the likelihood of transmission between any two plants to depend on the distance between them [19, 21, 26–28, 30]. Considering a spatially heterogeneous model would raise additional questions regarding the optimal spatial placement of sentinel plants and their selection as part of a monitoring programme, particularly if the risk of pathogen invasion also varied in space [65]. In addition to spatial heterogeneities, we could also consider temporal heterogeneities in the probability of invasion and detection that arise due to seasonal effects that impact vector dynamics [55, 66] and the level of symptoms displayed by infected hosts [36, 67, 68]. These heterogeneities motivate considering temporally varying sampling strategies that allow for sampling resources to be optimally allocated throughout the year. Further avenues for investigation include incorporating a latent period (time from when infection occurs to when the plant becomes infectious) in the model, or more generally a graduated progression through model compartments in which symptom expression and/or the likelihood of detection increase between successive compartments [69–71]. Similarly, we could alter our assumptions about the sensitivity and specificity of visual inspection as a method for detecting infection. For example, we could consider a sensitivity that increases over time as symptoms become more apparent, or allow for the possibility that detection occurs between sampling rounds.

In summary, our results represent a step towards understanding how sentinel plants may be used to facilitate the early detection of invasive plant pathogens. As we have shown, sentinels have the potential to reduce the expected incidence of disease upon pathogen discovery substantially, thereby increasing the chance of pathogen containment and lowering the cost of reactive control. These results encourage further research into the economic and logistical viability of using sentinel plants as a means of combating the problem of asymptomatic infection, as well as into the precise epidemiological characteristics of particular sentinel-crop-pathogen combinations. Monitoring programmes involving sentinel plants have the potential to reduce the negative impacts of a range of invading plant pathogens.

## Author contributions

Conceptualisation: All authors. Methodology: FALR, RNT. Investigation: FALR. Writing – original draft: FALR, RNT. Writing – review and editing: All authors. Supervision: RNT. All authors gave final approval for publication.

## Acknowledgements

The authors acknowledge the use of the University of Oxford Advanced Research Computing (ARC) facility which we used to run model simulations (http://dx.doi.org/10.5281/zenodo.22558). We also thank OJ Watson for early discussions about including sentinel plants in epidemiological models.

## Data accessibility

The computer code used to perform the analyses in this article is available in the following GitHub repository: https://github.com/francescalovellread/sentinel_plants. All computer code was written in the MATLAB programming environment (compatible with version R2022a).

## Ethics

This article does not present research with ethical considerations.

