## Supplementary text and figures for "Using ‘sentinel’ plants to improve early detection of invasive plant pathogens"

---

##### Text S1. Variation in parameter values

Where possible, the epidemiological parameters used in our main analyses were chosen based on literature estimates for *X. fastidiosa* infection in *O. europaea* and *C. roseus* (Table 1 of the main text). However, due to uncertainty in these reported values and the lack of available estimates of some parameters, we also conducted supplementary analyses to investigate the robustness of our results to parameter variation. The parameters that we varied, their meanings, their values taken in the main text and the alternative values considered in our supplementary analyses are given in Table S1. In almost all cases considered, the selected parameter was varied whilst all other parameters remained fixed at their main text values. The exception to this was when varying the crop population size ($P_C$): in those cases, we simultaneously scaled the transmission coefficients for 'Detectable' crops and sentinels ($\beta_C$ and $\beta_S$ respectively) to ensure that the epidemic growth rate remained fixed at $\beta_C P_C = 0.05$ [1]. For each alternative parameter value considered, we present plots analogous to Fig 6 in the main text, showing the optimal total number of sentinels in the population, the percentage change in EDP compared to the baseline level, the resultant EDP and the relative utility of a single sentinel plant (Supplementary Figs S1-14).

**Table S1. The parameters that we varied, their meanings, their values used in the main text and the alternative values we considered in our Supplementary analyses.**

| Parameter | Meaning | Main text value | Alternative values considered |
|---|---|---|---|
| $\beta_S$ | Transmission coefficient for 'Detectable' sentinels | $5 \times 10^{-5}$ | $2.5 \times 10^{-5}, 1 \times 10^{-4}$ |
| $\epsilon_C$ | Transmission scaling factor for 'Undetectable' crops | $0.015$ | $0.1, 0.25$ |
| $\epsilon_S$ | Transmission scaling factor for 'Undetectable' sentinels | $0.1$ | $0.02, 0.5$ |
| $\gamma_C$ | Duration of crop 'Undetectable' period | $452$ days | $350$ days, $550$ days |
| $\gamma_S$ | Duration of sentinel 'Undetectable' period | $49$ days | $28$ days, $70$ days |
| $P_C$ | Number of crop plants in the population | $1000$ | $500, 1500$* |
| $U_0$ | Initial number of 'Undetectable' infected individuals | $1$ | $2, 4$ |

* values of $\beta_C, \beta_S$ scaled accordingly to ensure $\beta_C P_C = 0.05$.

### Text S2. Bayesian optimisation

Overview of Bayesian optimisation

Bayesian optimisation is an efficient technique for finding the extrema of objective functions that are expensive to evaluate [2]. It is an iterative process in which successive (possibly noisy) observations of an objective function are taken at carefully selected trial points and used to update our beliefs about its likeliest form. Beginning from a prior distribution on the objective function (invariably a Gaussian process prior [2, 3]), on each iteration the objective function is sampled at a trial point and Bayes' theorem is applied to incorporate the observed result into a posterior distribution. The extrema of this posterior approximate those of the objective function with increasing accuracy on each iteration.

Successive trial points are selected by maximising an acquisition function that quantifies the potential gain of sampling at a particular point. When seeking to minimise the objective function, the acquisition function is typically defined so that it takes high values i) near to previously sampled points where the objective function is small (known as exploitation), and ii) where there is a large amount of uncertainty, and the objective function has the potential to be small (known as exploration). In contrast to the objective function, the acquisition function is cheap to evaluate and maximise, so the best trial point can be found easily.

The judicious selection of trial points based on their utility makes Bayesian optimisation a particularly efficient method in terms of the number of evaluations of the objective function required, which is very useful when sampling is expensive [2]. Furthermore, Bayesian optimisation can be applied to an objective function that does not have a known analytic form, as long as the function can be evaluated computationally for a given choice of input parameters. It is therefore an ideal optimisation technique for this study, since our objective function is both analytically inaccessible and expensive to evaluate computationally.

Application of Bayesian optimisation in this study

In this study, we implemented Bayesian optimisation in MATLAB using the inbuilt function '*bayesopt*' [4, 5]. Since '*bayesopt*' seeks to minimise (rather than maximise) a specified objective function, we defined our objective to be the percentage change in EDP compared to the baseline level for a given $(N, \Delta)$ pair. Therefore, smaller (more negative) values of the objective corresponded to greater reductions in the EDP.

Initially, we fixed the number of sentinels added to the population ($P_S$) and considered the number of sentinels included in the sample ($N_S$) as the single optimisable variable, allowed to vary between 0 and the (constant) upper bound $\min(P_S, N)$ (see Section 3.2 of the main text). We used the '*expected-improvement-plus*' acquisition function, which selects the next trial point (in this case value of $N_S$) so as to maximise the expected reduction in the objective function compared to the current estimated minimum, whilst also avoiding overexploitation of any given area [4]. For each choice of $(N, \Delta)$ we performed 30 iterations of the Bayesian optimisation algorithm (30 iterations is the *bayesopt* default); that is, we evaluated the objective function for 30 successively selected values of $N_S$ (not necessarily distinct). Each evaluation of the objective function involved computing the resultant EDP for the sampling strategy under consideration by performing sampling on 25,000 simulated epidemic curves (as shown in Fig 2 of the main text), and comparing it to the baseline EDP for the specified $N$ and $\Delta$.

Subsequently, we allowed both the number of sentinels added to the population ($P_S$) and the number of sentinels included in the sample ($N_S$) to vary simultaneously, and adapted the Bayesian optimisation algorithm to treat each of these as an optimisable variable (see Section 3.3 of the main text). We allowed $P_S$ to vary between 0 and 350, and $N_S$ to vary between 0 and $\min(P_S, N)$ (now variable). Since the default implementation of '*bayesopt*' requires constant bounds for the optimisable variables, we first specified that $N_S$ could vary between 0 and the total sample size $N$. Then, we further restricted the feasible region from which trial points could be drawn by enforcing the constraint $N_S \leq P_S$. This form of constraint (a deterministic function of the optimisable variables) can be implemented in MATLAB using the '*XConstraintFcn*' option in '*bayesopt*' [6].

Applying Bayesian optimisation in this study resulted in a substantial reduction in the number of objective function evaluations required, and therefore a much shorter computation time. For example, in Fig 6 we considered 900 distinct $(N, \Delta)$ pairs. For each of these pairs, $P_S$ was allowed to vary in the range $[0{,}350]$, and $N_S$ in the range $\min(P_S, N)$. Seeking the optimal combination of $P_S$ and $N_S$ by exhaustive search would have required tens of thousands of objective function evaluations for each $(N, \Delta)$ pair; by applying Bayesian optimisation, this was reduced to 30 iterations per pair.

#### Text S3. Random sampling vs. repeated sampling

When considering the use of sentinel plants for surveillance, as well as deciding how many sentinels to add to the population ($P_S$) an important question is how many of those sentinels to include in the sample ($N_S$). A natural choice is to sample the maximum possible number of sentinels – that is, choosing $N_S = \min(P_S, N)$. However, in some circumstances this would result in frequent resampling of the same plants. For example, if the total sample size $N$ is equal to the total number of sentinels in the population $P_S$, selecting $N_S$ in this way would mean inspecting an identical set of plants on every sampling round. Since the information we gain about the disease status of a plant we have already inspected in the recent past is less than the information we gain by looking at a plant we have not previously inspected, repeated sampling of this kind may reduce the effectiveness of the surveillance strategy compared to a situation in which the sample selection has an element of randomness.

Here, we use a simple example to demonstrate how repeated sampling of the same plants can lead to a worse outcome than selecting plants to sample at random. We constructed a standard one-species Susceptible-Infected (SI) model in which plants progress from the 'Susceptible' class to the 'Infected' class at rate $\beta SI$ (where $\beta$ is the transmission coefficient, $S$ is the number of 'Susceptible' plants and $I$ is the number of 'Infected' plants). Setting $\beta = 5 \times 10^{-6}$, we generated simulated epidemic curves in a population of size $1000$, beginning from a single infected individual. We considered a monitoring strategy in which samples of size $N$ were taken from the population every $\Delta$ days, with detection occurring if and only if an 'Infected' plant was included in the sample. We implemented this monitoring strategy on our simulated epidemic curves for a range of sample sizes $N$ and sample intervals $\Delta$. For each choice of $N$ and $\Delta$ we considered both random sampling (in which the sampled plants were chosen at random on every sampling round), and repeated sampling (in which the same $N$ plants were inspected on every sampling round).

For the chosen model parameterisation, random sampling outperformed repeated sampling across the range of $(N, \Delta)$ values considered (Supplementary Fig S15). This result is conceptually equivalent to the phenomenon observed in the main text – that sampling as many sentinels as possible is not always the best strategy, if repeated sampling is required to achieve this. Note that the difference between the two sampling approaches was smaller for large sample intervals $\Delta$, since the longer the time between samples the less relevant the information gained from a previous sample is to the current state of the system. Therefore, less information is lost by looking at the same plants. The difference between the two approaches also decreased for larger samples sizes $N$, since detection occurred within fewer sampling rounds (thus less repetition occurred).

#### Text S4. Variation in the objective function

Throughout our main analyses, we investigated how including sentinel plants in a surveillance strategy could reduce the EDP compared to the baseline level, where the EDP was defined to be the expected detection prevalence in the crop population at the time of discovery. In doing so, we assumed that minimising the prevalence of infection amongst valuable crop plants was likely to be the primary objective of the grower, and did not consider the prevalence of infection in the sentinel population. However, in practice a grower may also wish to take the expected detection prevalence in the sentinel population ($EDP_{sent}$) into account in some way when evaluating the effects of surveillance strategies. Reasons for this include reducing the cost of removing infected plants post-detection and reducing the likelihood that the pathogen is exported to a new location before control can be implemented. Additionally, in some pathosystems the sentinel plants may have some intrinsic value – for example, young orange trees used as sentinels for greening disease in citrus groves (see Discussion).

Our analyses can be adapted to explore any balance of interest between crop and sentinel discovery prevalence by replacing the EDP with $\Omega = \text{EDP} + (\omega_S \times \text{EDP}_{\text{sent}})$ for some weight $\omega_S \in [0,1]$. Computationally speaking, this simply requires a small adjustment to the objective function used in our Bayesian optimisation algorithm. Our main analyses correspond to the case $\omega_S = 0$ (Fig 6): here, we display analogous results for the case $\omega_S = 0.5$ (Supplementary Fig S18). This represents a hypothetical situation in which an infected sentinel plant is considered half as concerning as an infected crop plant (rather than not being considered at all). In this case, introducing a large number of sentinel plants is penalised by contributing to an increased $\text{EDP}_{\text{sent}}$. The optimal number of sentinels to include in the population is therefore reduced compared to the case in which $\omega_S = 0$, apart from when the sample interval $\Delta$ is large and the optimal number of sentinels is equal to the sample size (i.e. not in excess) (Supplementary Fig S18A). The achievable percentage reduction in $\Omega$ compared to the baseline level is also reduced compared to the case in which we consider solely the EDP, since infected sentinel plants are no longer discounted (Supplementary Fig S18B). This confirms that the specific objective a grower wishes to achieve should be carefully considered before the results of sentinel-based surveillance strategies are evaluated.

#### *Supplementary Figures*

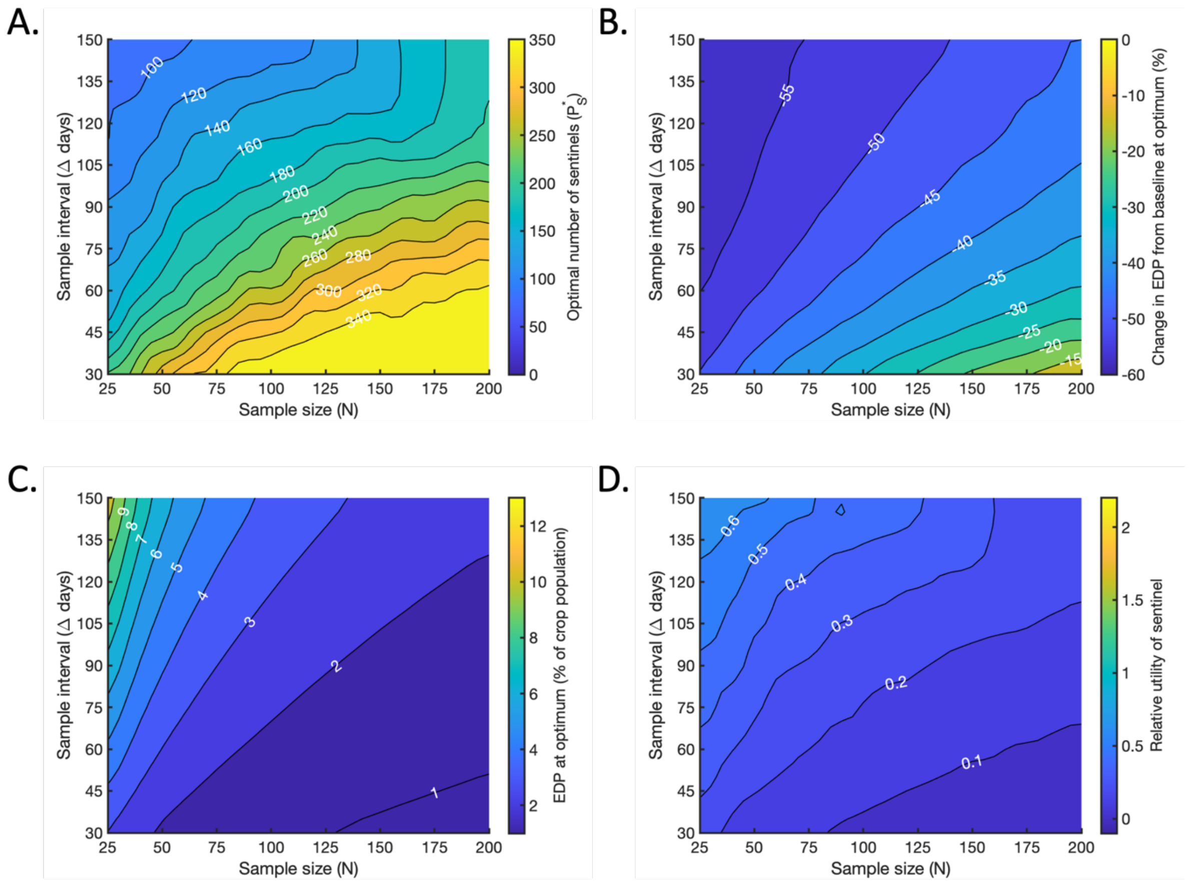


**Fig S1. The effect of decreasing the transmission coefficient for 'Detectable' sentinels from $\beta_S = 5 \times 10^{-5}$ (baseline value) to $\beta_S = 2.5 \times 10^{-5}$. Panels analogous to Fig 6 in the main text.** A. The optimal number $P_S^*$ of sentinel plants to include in the population, for which the maximal reduction in the EDP compared to the baseline level is achieved (if $N_S$ is also chosen optimally). B. The percentage change in the EDP compared to the baseline value at the optimum, achieved when $P_S = P_S^*$ and $N_S = N_S^*$. C. The resultant value of the EDP at the optimum, expressed as a percentage of the total crop population. D. The relative utility of a single sentinel plant (the percentage reduction in EDP per sentinel) at the optimum.

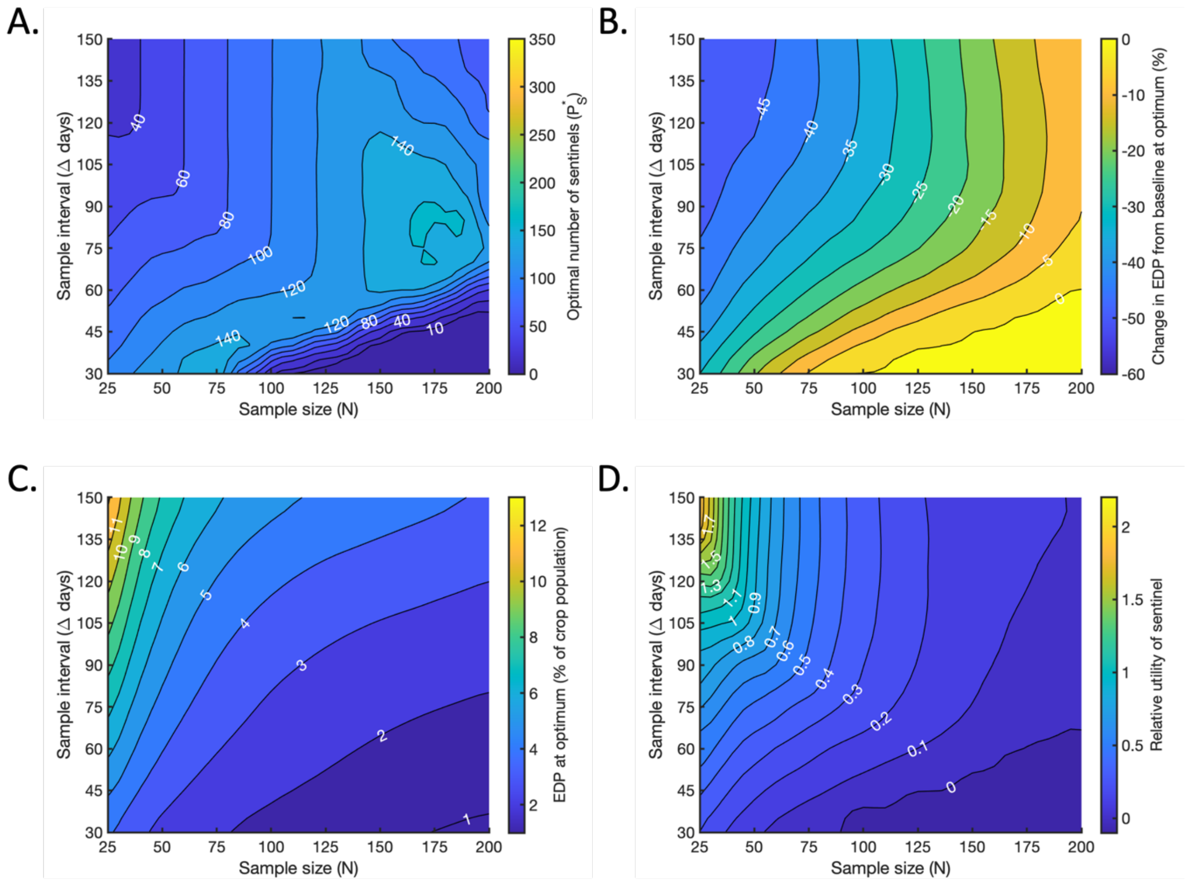


**Fig S2. The effect of increasing the transmission coefficient for 'Detectable' sentinels from $\boldsymbol{\beta_S} = \mathbf{5 \times 10^{-5}}$ (baseline value) to $\boldsymbol{\beta_S} = \mathbf{1 \times 10^{-4}}$. Panels analogous to Fig 6 in the main text.** A. The optimal number $P_S^*$ of sentinel plants to include in the population, for which the maximal reduction in the EDP compared to the baseline level is achieved (if $N_S$ is also chosen optimally). B. The percentage change in the EDP compared to the baseline value at the optimum, achieved when $P_S = P_S^*$ and $N_S = N_S^*$. C. The resultant value of the EDP at the optimum, expressed as a percentage of the total crop population. D. The relative utility of a single sentinel plant (the percentage reduction in EDP per sentinel) at the optimum.

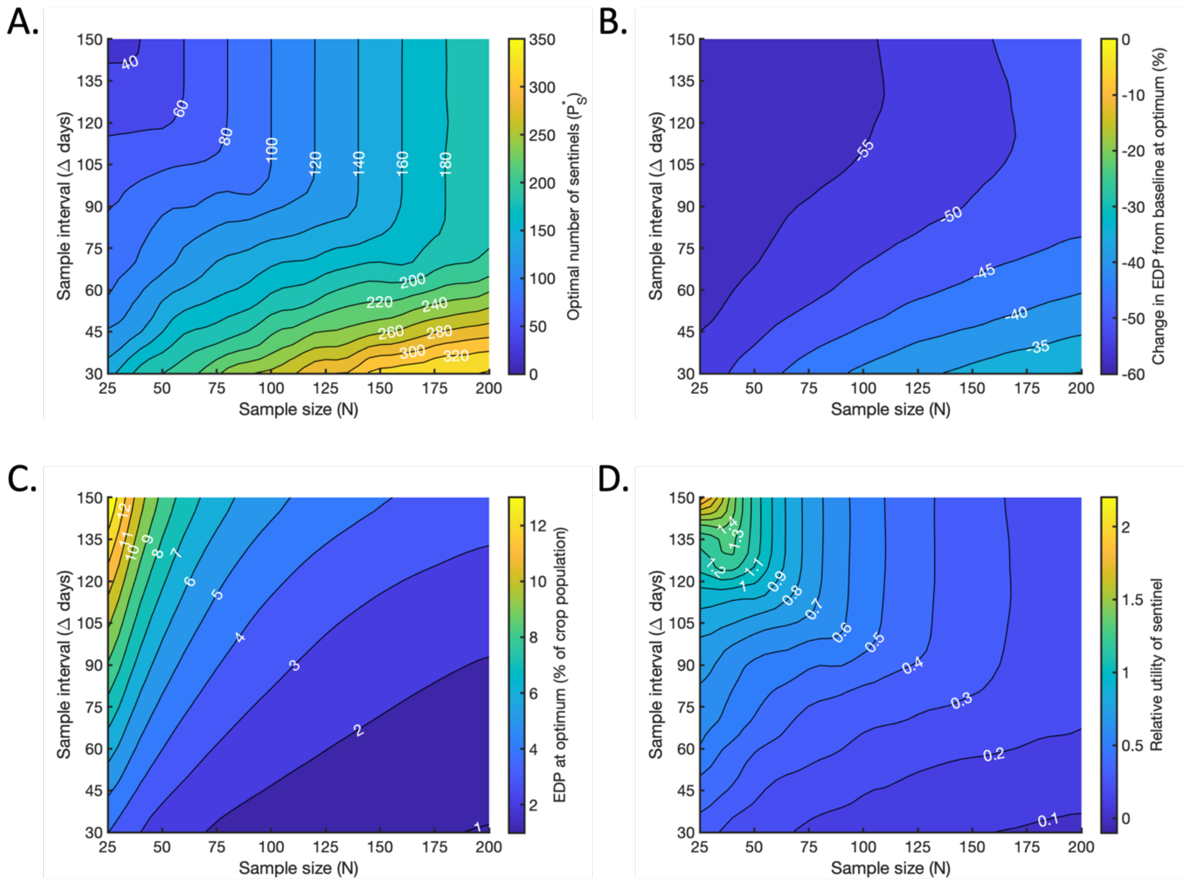


**Fig S3. The effect of increasing the transmission scaling factor for 'Undetectable' crops from $\epsilon_C = 0.015$ (baseline value) to $\epsilon_C = 0.1$. Panels analogous to Fig 6 in the main text.** A. The optimal number $P_S^*$ of sentinel plants to include in the population, for which the maximal reduction in the EDP compared to the baseline level is achieved (if $N_S$ is also chosen optimally). B. The percentage change in the EDP compared to the baseline value at the optimum, achieved when $P_S = P_S^*$ and $N_S = N_S^*$. C. The resultant value of the EDP at the optimum, expressed as a percentage of the total crop population. D. The relative utility of a single sentinel plant (the percentage reduction in EDP per sentinel) at the optimum.

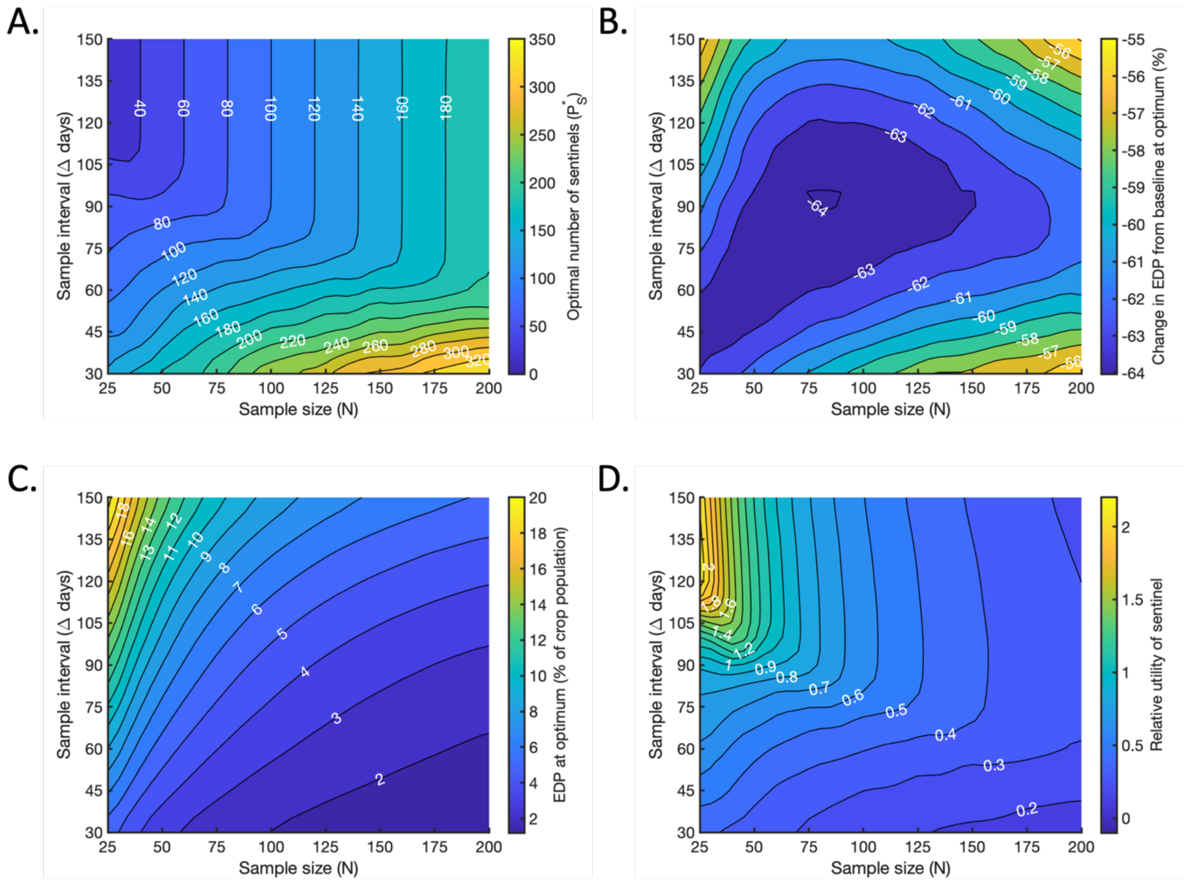


**Fig S4. The effect of increasing the transmission scaling factor for 'Undetectable' crops from $\epsilon_C = 0.015$ (baseline value) to $\epsilon_C = 0.25$. Panels analogous to Fig 6 in the main text.** A. The optimal number $P_S^*$ of sentinel plants to include in the population, for which the maximal reduction in the EDP compared to the baseline level is achieved (if $N_S$ is also chosen optimally). B. The percentage change in the EDP compared to the baseline value at the optimum, achieved when $P_S = P_S^*$ and $N_S = N_S^*$. C. The resultant value of the EDP at the optimum, expressed as a percentage of the total crop population. D. The relative utility of a single sentinel plant (the percentage reduction in EDP per sentinel) at the optimum.

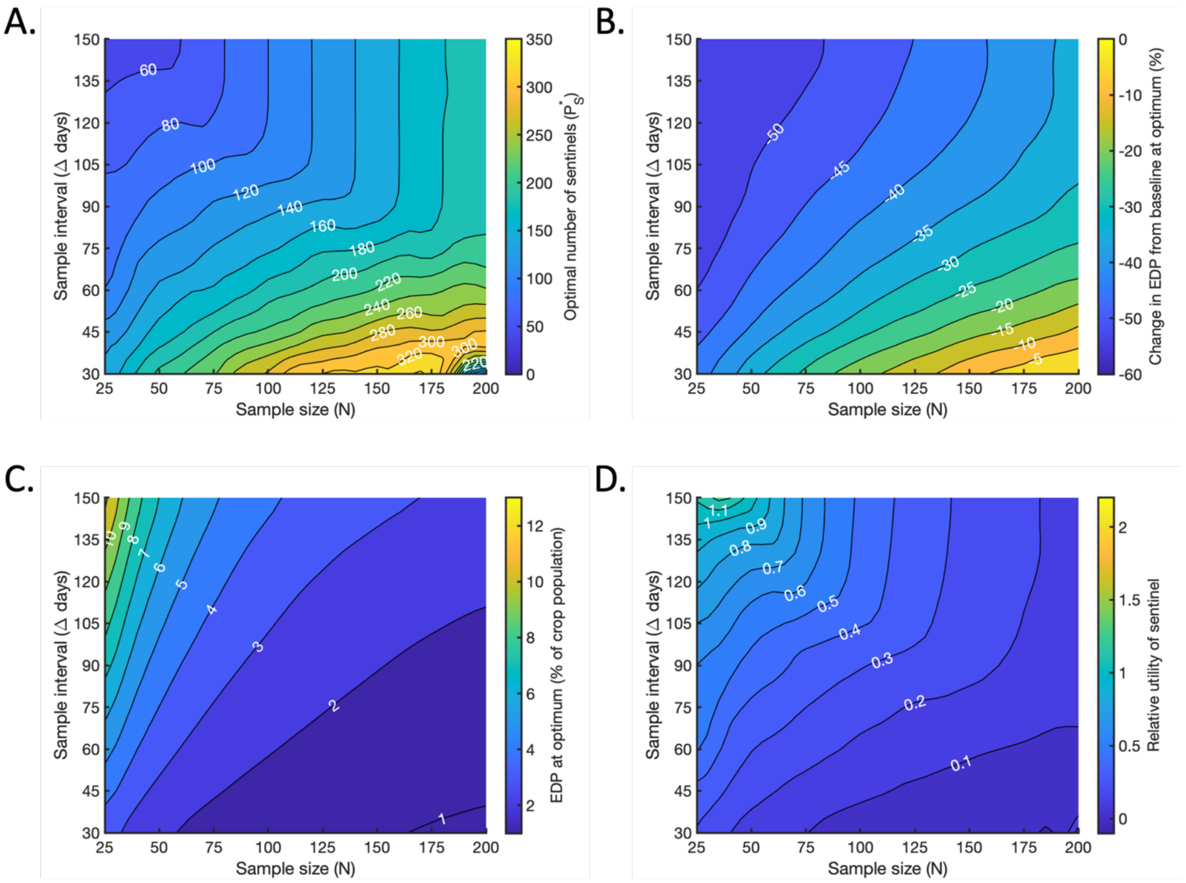


**Fig S5. The effect of reducing the transmission scaling factor for 'Undetectable' sentinels from $\epsilon_S = 0.1$ (baseline value) to $\epsilon_S = 0.02$. Panels analogous to Fig 6 in the main text.** A. The optimal number $P_S^*$ of sentinel plants to include in the population, for which the maximal reduction in the EDP compared to the baseline level is achieved (if $N_S$ is also chosen optimally). B. The percentage change in the EDP compared to the baseline value at the optimum, achieved when $P_S = P_S^*$ and $N_S = N_S^*$. C. The resultant value of the EDP at the optimum, expressed as a percentage of the total crop population. D. The relative utility of a single sentinel plant (the percentage reduction in EDP per sentinel) at the optimum.

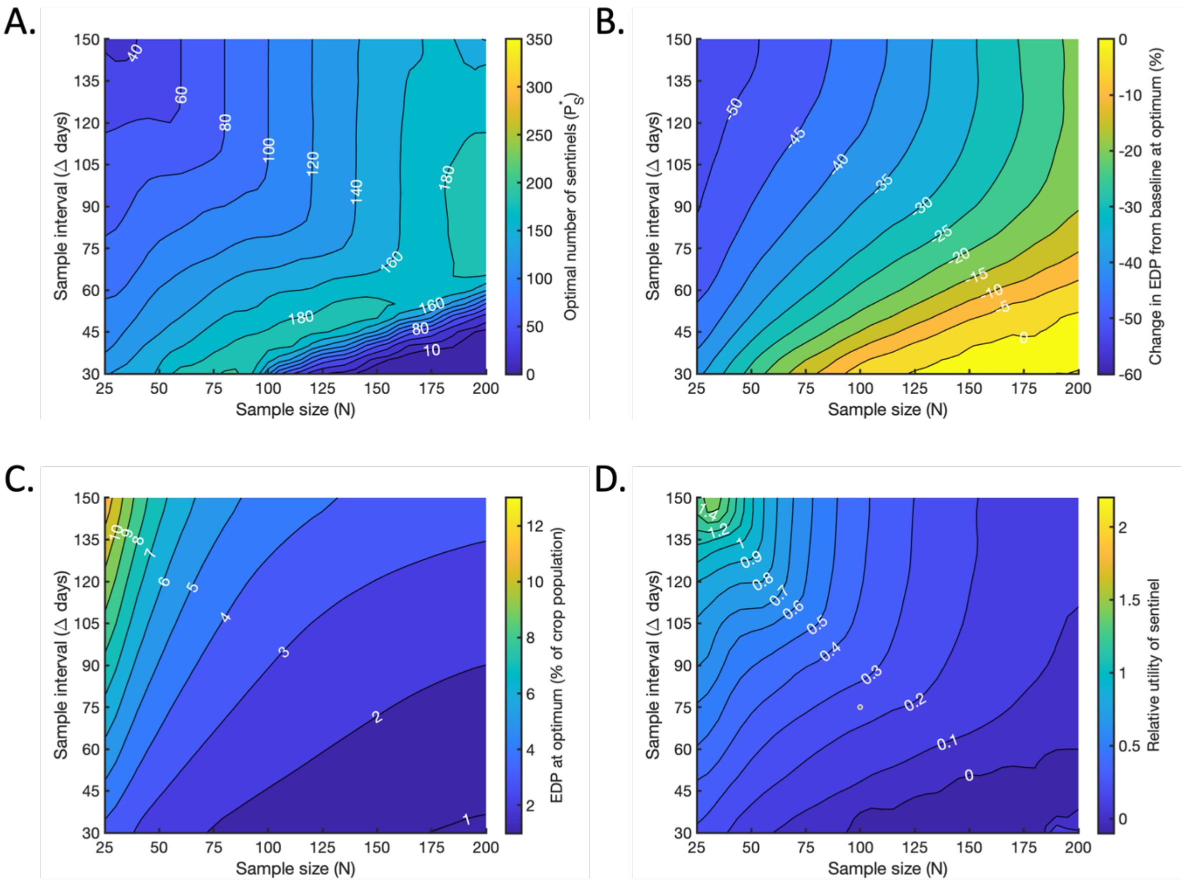


**Fig S6. The effect of increasing the transmission scaling factor for 'Undetectable' sentinels from $\epsilon_S = 0.1$ (baseline value) to $\epsilon_S = 0.5$. Panels analogous to Fig 6 in the main text.** A. The optimal number $P_S^*$ of sentinel plants to include in the population, for which the maximal reduction in the EDP compared to the baseline level is achieved (if $N_S$ is also chosen optimally). B. The percentage change in the EDP compared to the baseline value at the optimum, achieved when $P_S = P_S^*$ and $N_S = N_S^*$. C. The resultant value of the EDP at the optimum, expressed as a percentage of the total crop population. D. The relative utility of a single sentinel plant (the percentage reduction in EDP per sentinel) at the optimum.

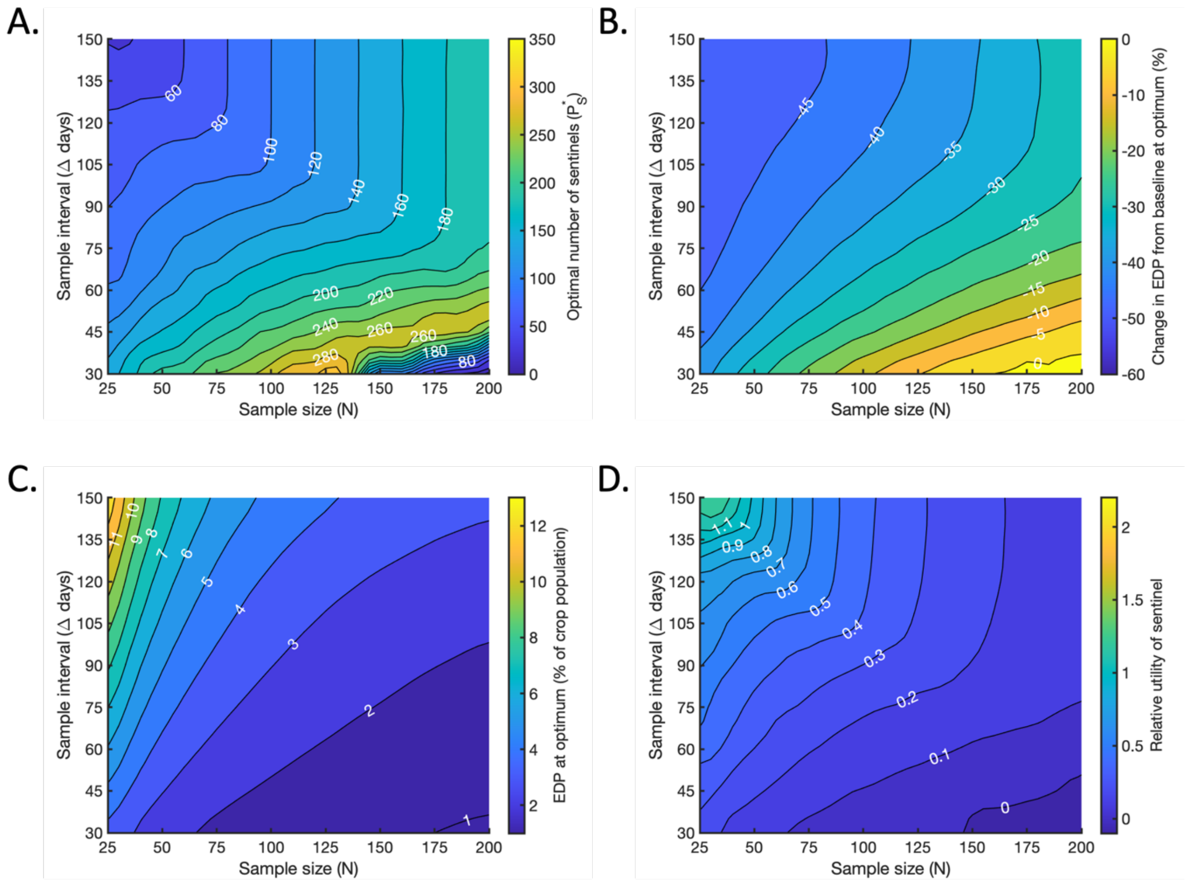


**Fig S7. The effect of reducing the duration of the crop 'Undetectable' period from $\gamma_C = 452$ days (baseline value) to $\gamma_C = 350$ days. Panels analogous to Fig 6 in the main text.** A. The optimal number $P_S^*$ of sentinel plants to include in the population, for which the maximal reduction in the EDP compared to the baseline level is achieved (if $N_S$ is also chosen optimally). B. The percentage change in the EDP compared to the baseline value at the optimum, achieved when $P_S = P_S^*$ and $N_S = N_S^*$. C. The resultant value of the EDP at the optimum, expressed as a percentage of the total crop population. D. The relative utility of a single sentinel plant (the percentage reduction in EDP per sentinel) at the optimum.

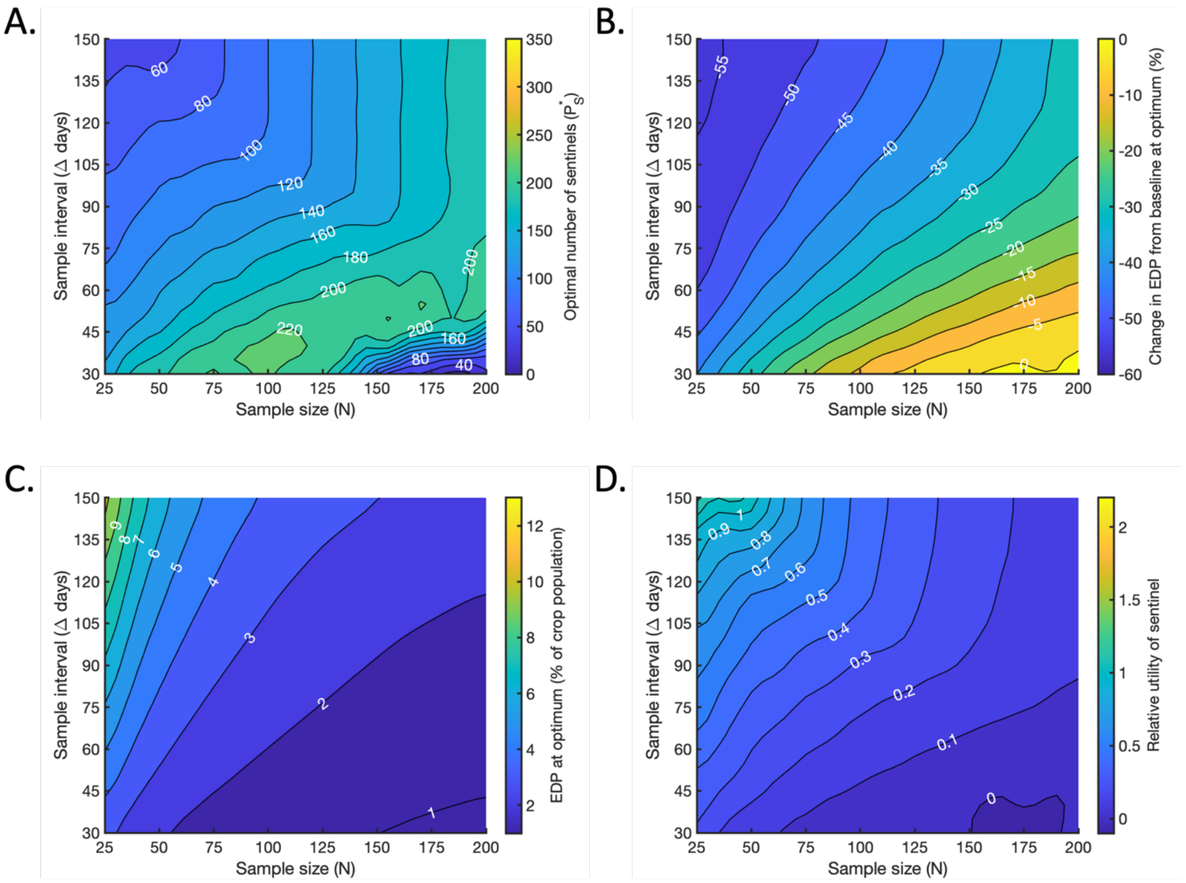


**Fig S8. The effect of increasing the duration of the crop 'Undetectable' period from $\gamma_C = 452$ days (baseline value) to $\gamma_C = 550$ days. Panels analogous to Fig 6 in the main text.** A. The optimal number $P_S^*$ of sentinel plants to include in the population, for which the maximal reduction in the EDP compared to the baseline level is achieved (if $N_S$ is also chosen optimally). B. The percentage change in the EDP compared to the baseline value at the optimum, achieved when $P_S = P_S^*$ and $N_S = N_S^*$. C. The resultant value of the EDP at the optimum, expressed as a percentage of the total crop population. D. The relative utility of a single sentinel plant (the percentage reduction in EDP per sentinel) at the optimum.

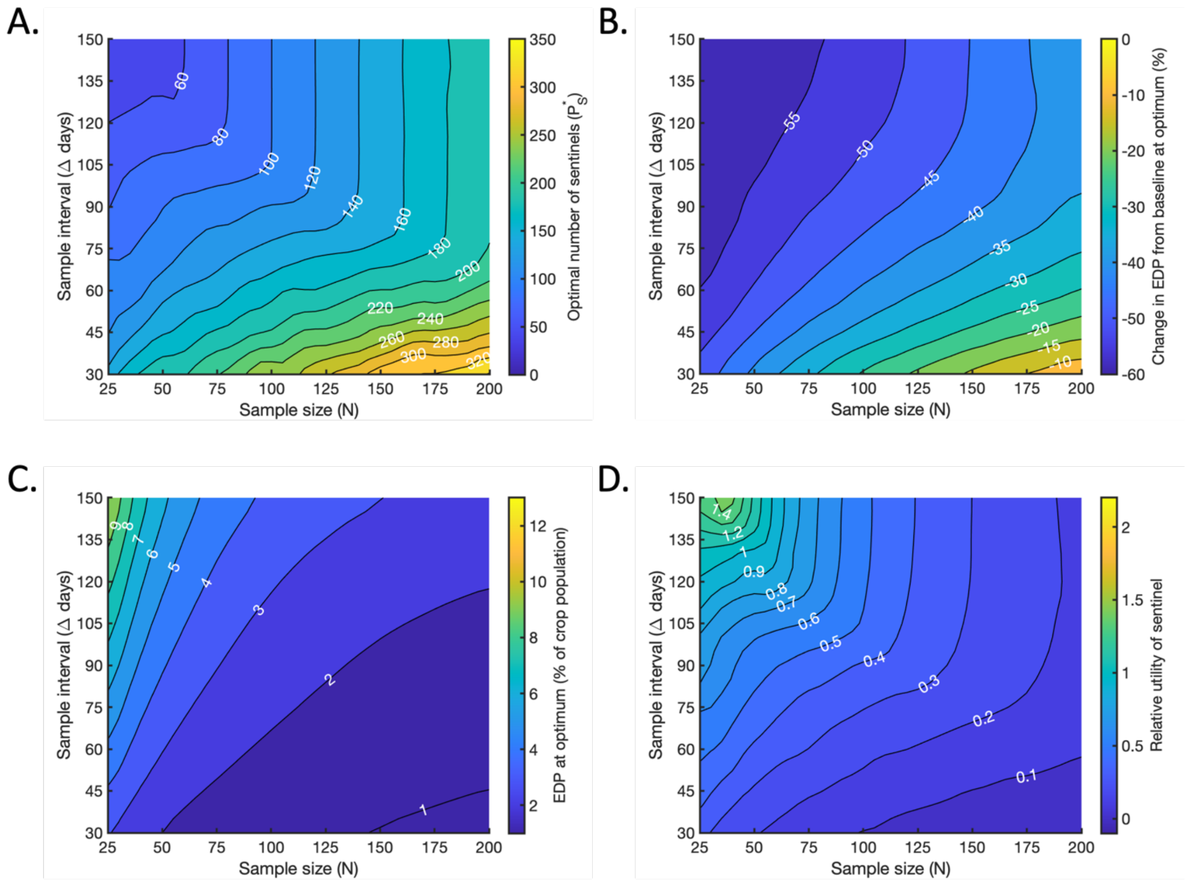


**Fig S9. The effect of reducing the duration of the sentinel 'Undetectable' period from $\gamma_S = 49$ days (baseline value) to $\gamma_S = 28$ days. Panels analogous to Fig 6 in the main text.** A. The optimal number $P_S^*$ of sentinel plants to include in the population, for which the maximal reduction in the EDP compared to the baseline level is achieved (if $N_S$ is also chosen optimally). B. The percentage change in the EDP compared to the baseline value at the optimum, achieved when $P_S = P_S^*$ and $N_S = N_S^*$. C. The resultant value of the EDP at the optimum, expressed as a percentage of the total crop population. D. The relative utility of a single sentinel plant (the percentage reduction in EDP per sentinel) at the optimum.

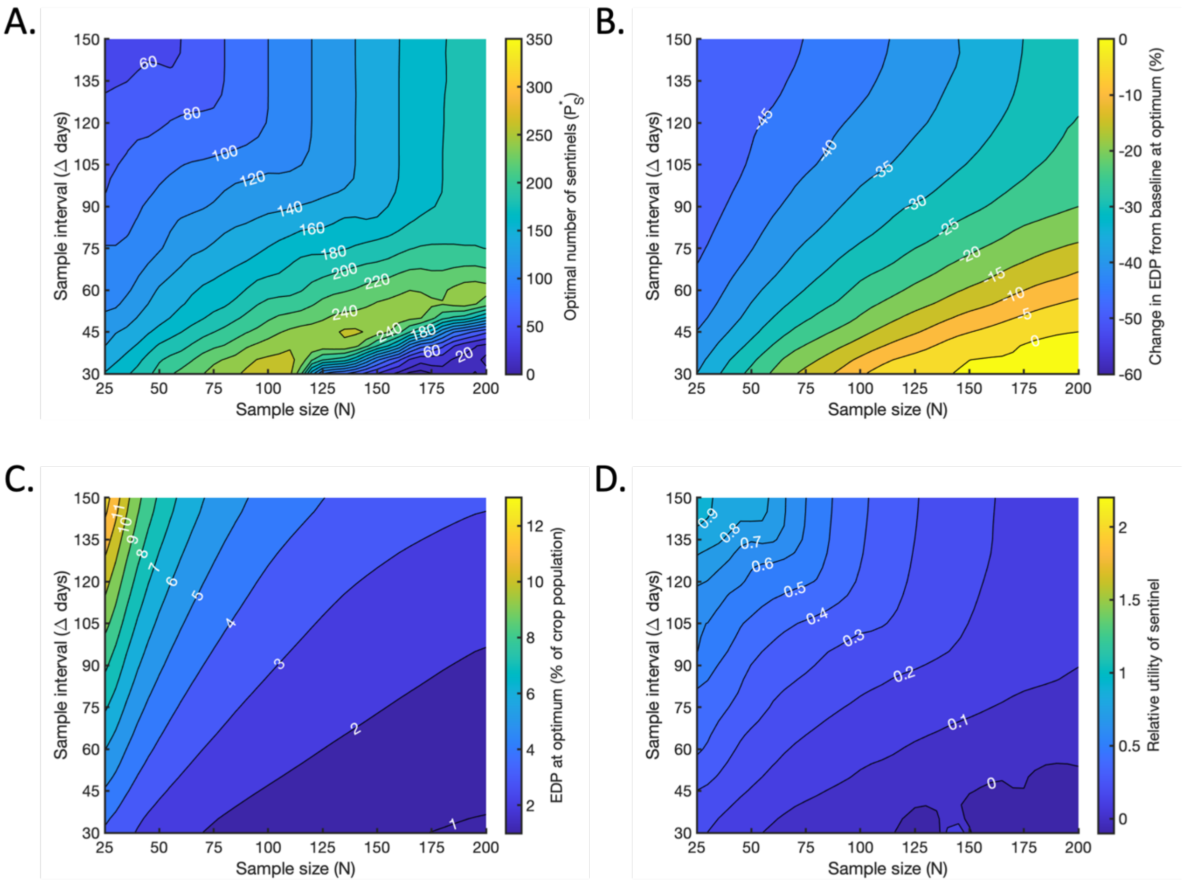


**Fig S10. The effect of increasing the duration of the sentinel 'Undetectable' period from $\gamma_S = 49$ days (baseline value) to $\gamma_S = 70$ days. Panels analogous to Fig 6 in the main text.** A. The optimal number $P_S^*$ of sentinel plants to include in the population, for which the maximal reduction in the EDP compared to the baseline level is achieved (if $N_S$ is also chosen optimally). B. The percentage change in the EDP compared to the baseline value at the optimum, achieved when $P_S = P_S^*$ and $N_S = N_S^*$. C. The resultant value of the EDP at the optimum, expressed as a percentage of the total crop population. D. The relative utility of a single sentinel plant (the percentage reduction in EDP per sentinel) at the optimum.

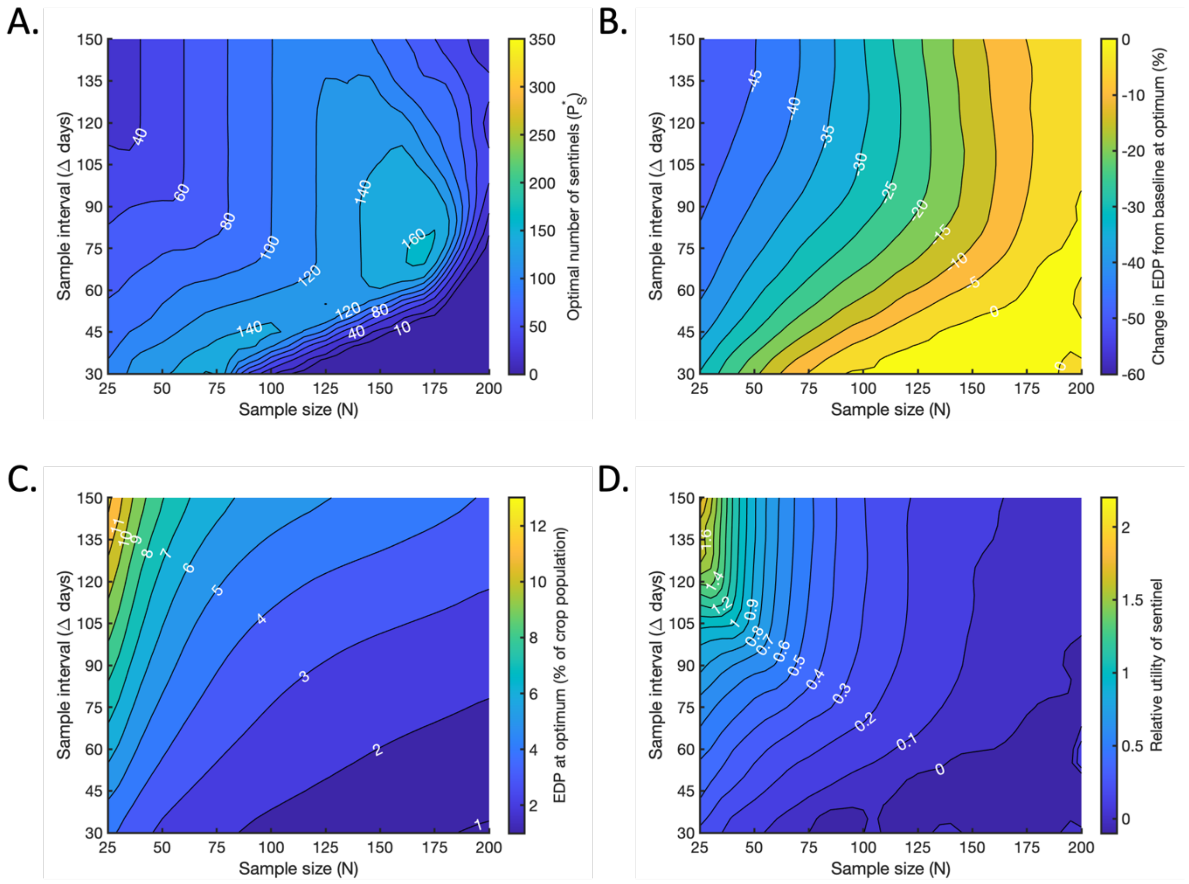


**Fig S11. The effect of decreasing the crop population size from $P_C = 1000$ plants (baseline value) to $P_C = 500$ plants. Panels analogous to Fig 6 in the main text.** A. The optimal number $P_S^*$ of sentinel plants to include in the population, for which the maximal reduction in the EDP compared to the baseline level is achieved (if $N_S$ is also chosen optimally). B. The percentage change in the EDP compared to the baseline value at the optimum, achieved when $P_S = P_S^*$ and $N_S = N_S^*$. C. The resultant value of the EDP at the optimum, expressed as a percentage of the total crop population. D. The relative utility of a single sentinel plant (the percentage reduction in EDP per sentinel) at the optimum.

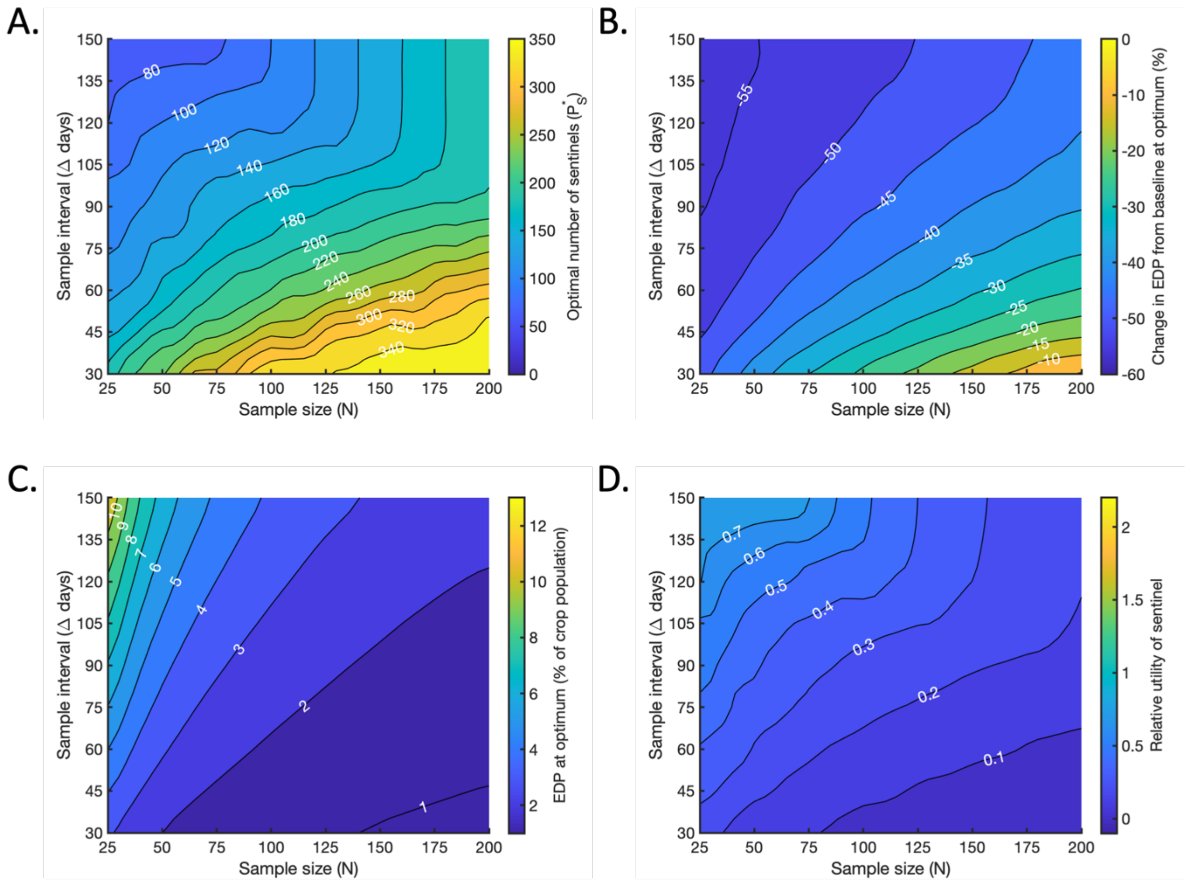


**Fig S12. The effect of increasing the crop population size from $P_C = 1000$ plants (baseline value) to $P_C = 1500$ plants. Panels analogous to Fig 6 in the main text.** A. The optimal number $P_S^*$ of sentinel plants to include in the population, for which the maximal reduction in the EDP compared to the baseline level is achieved (if $N_S$ is also chosen optimally). B. The percentage change in the EDP compared to the baseline value at the optimum, achieved when $P_S = P_S^*$ and $N_S = N_S^*$. C. The resultant value of the EDP at the optimum, expressed as a percentage of the total crop population. D. The relative utility of a single sentinel plant (the percentage reduction in EDP per sentinel) at the optimum.

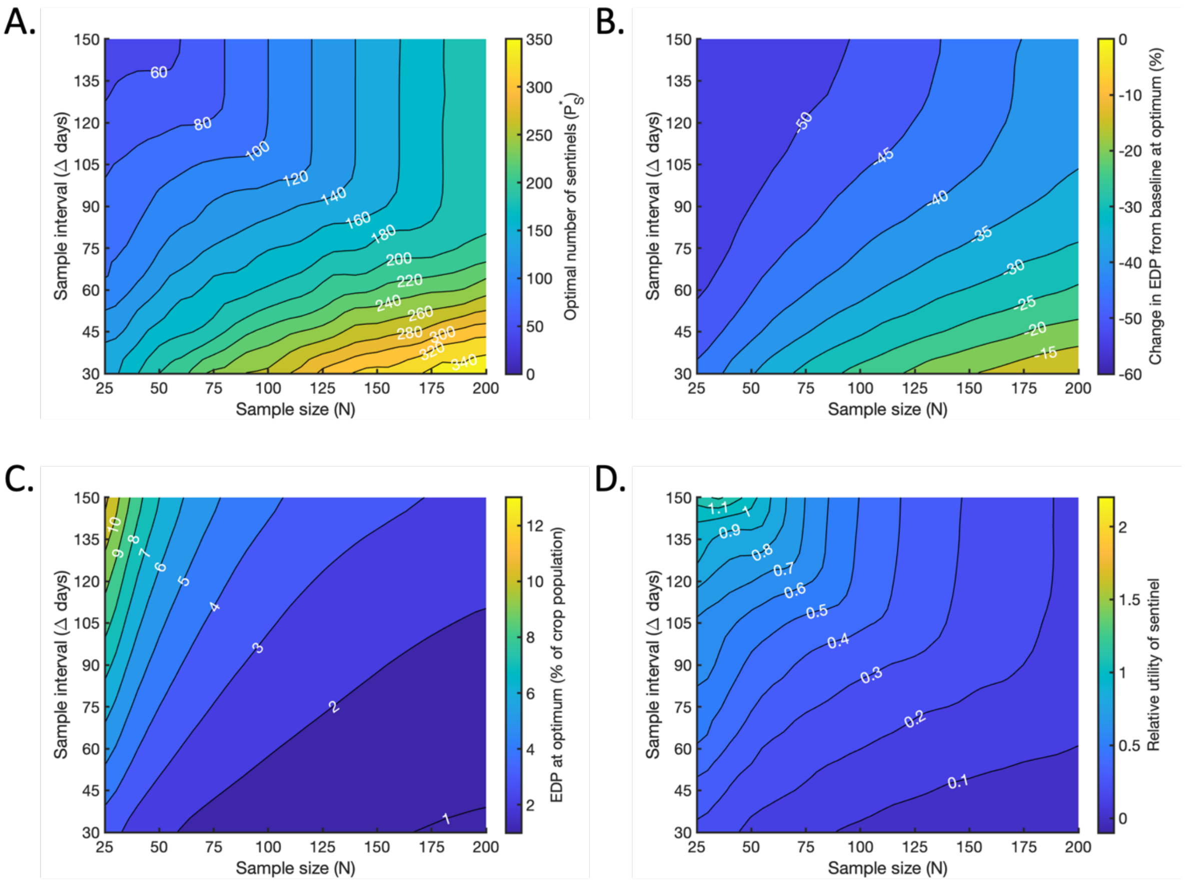


**Fig S13. The effect of increasing the initial number of infected plants from $U_0 = 1$ (baseline value) to $U_0 = 2$. Panels analogous to Fig 6 in the main text.** A. The optimal number $P_S^*$ of sentinel plants to include in the population, for which the maximal reduction in the EDP compared to the baseline level is achieved (if $N_S$ is also chosen optimally). B. The percentage change in the EDP compared to the baseline value at the optimum, achieved when $P_S = P_S^*$ and $N_S = N_S^*$. C. The resultant value of the EDP at the optimum, expressed as a percentage of the total crop population. D. The relative utility of a single sentinel plant (the percentage reduction in EDP per sentinel) at the optimum.

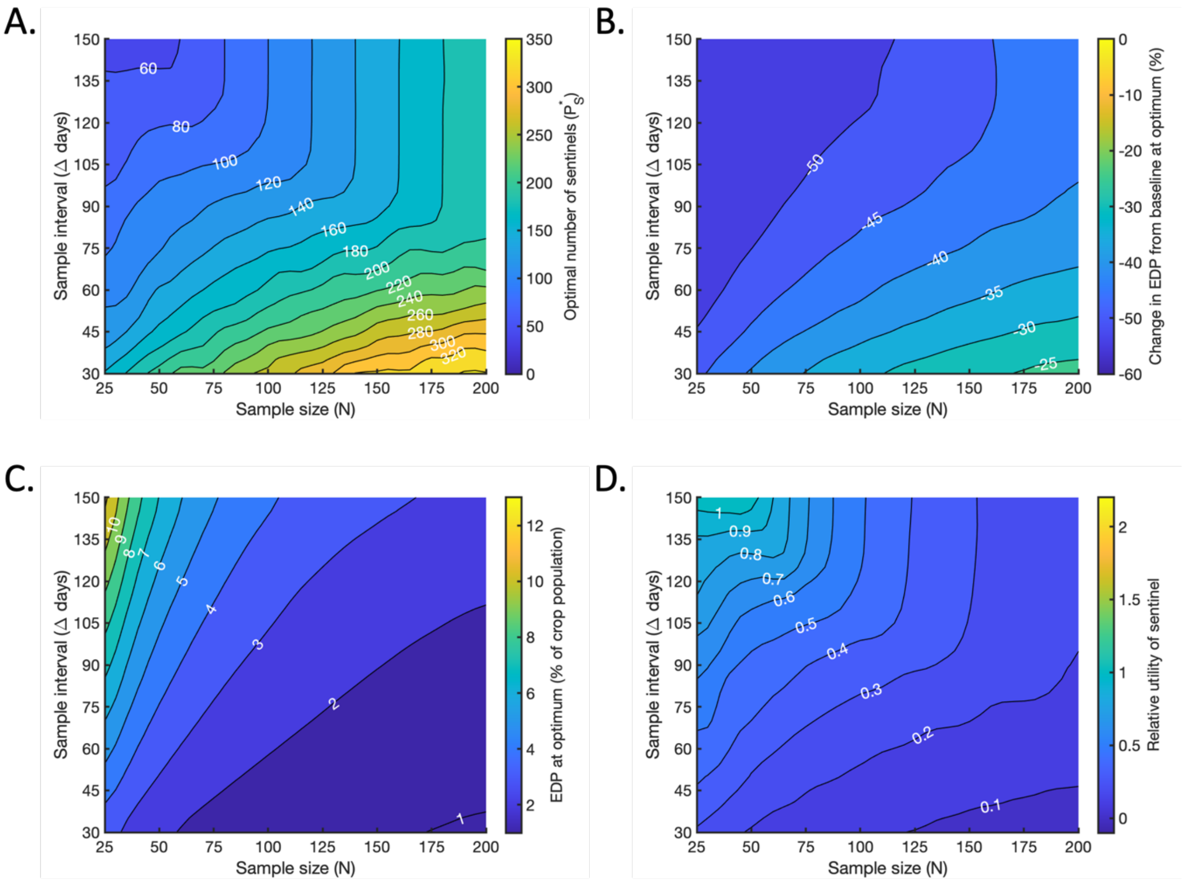


**Fig S14. The effect of increasing the initial number of infected plants from $U_0 = 1$ (baseline value) to $U_0 = 4$. Panels analogous to Fig 6 in the main text.** A. The optimal number $P_S^*$ of sentinel plants to include in the population, for which the maximal reduction in the EDP compared to the baseline level is achieved (if $N_S$ is also chosen optimally). B. The percentage change in the EDP compared to the baseline value at the optimum, achieved when $P_S = P_S^*$ and $N_S = N_S^*$. C. The resultant value of the EDP at the optimum, expressed as a percentage of the total crop population. D. The relative utility of a single sentinel plant (the percentage reduction in EDP per sentinel) at the optimum.

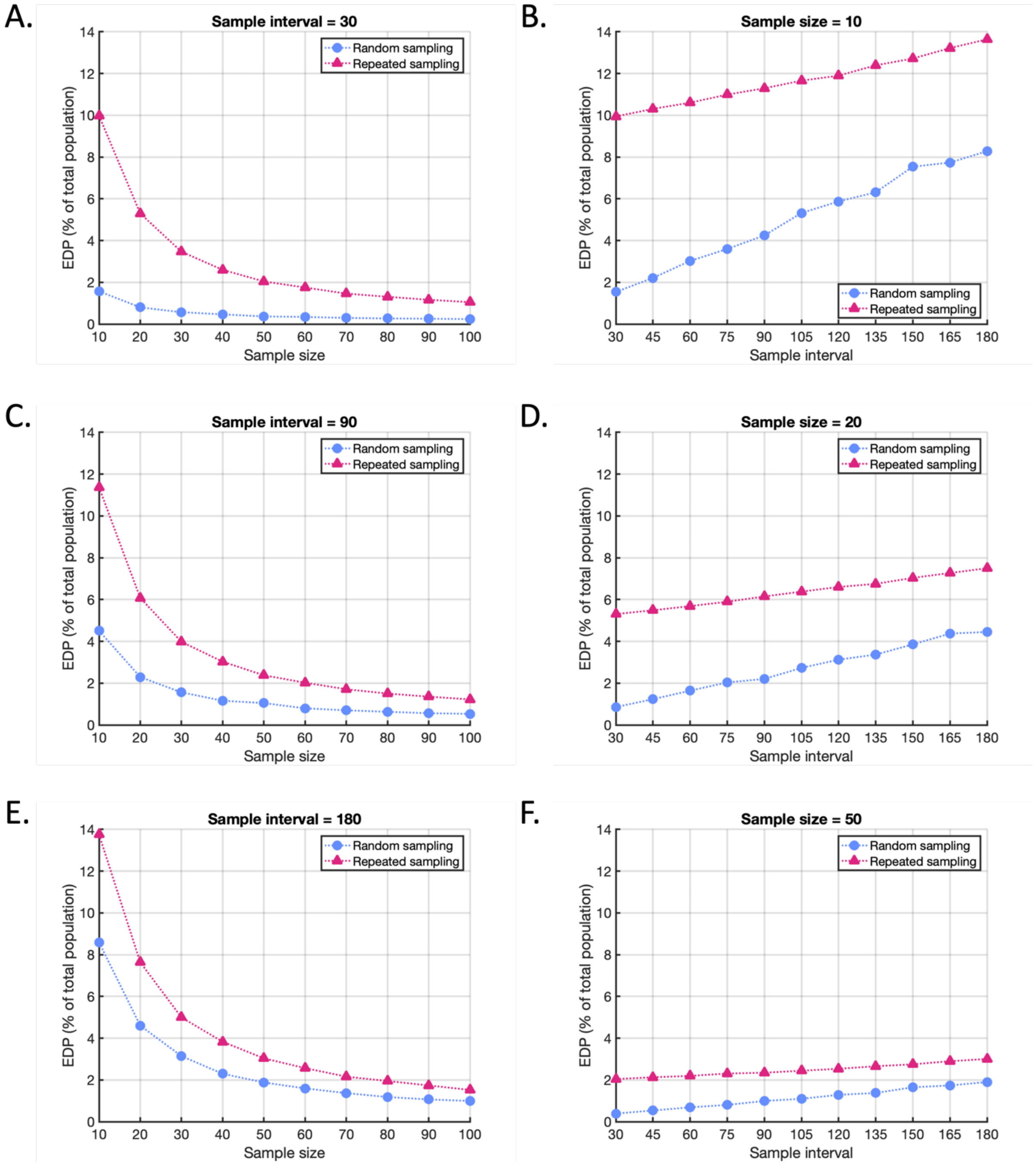


**Fig S15. Resultant EDPs (as a percentage of the total population) for the example system described in Supplementary Text S3, illustrating how random sampling (blue circles) outperforms repeated sampling (pink triangles).** A,C,E. The resultant EDP for sample intervals $\Delta = 30$ days (A), 90 days (C) and 180 days (E) as the sample size $N$ varies. B,D,F. The resultant EDP for sample sizes $N = 10$ (B), $N = 20$ (D) and $N = 50$ (F) as the sample interval varies.

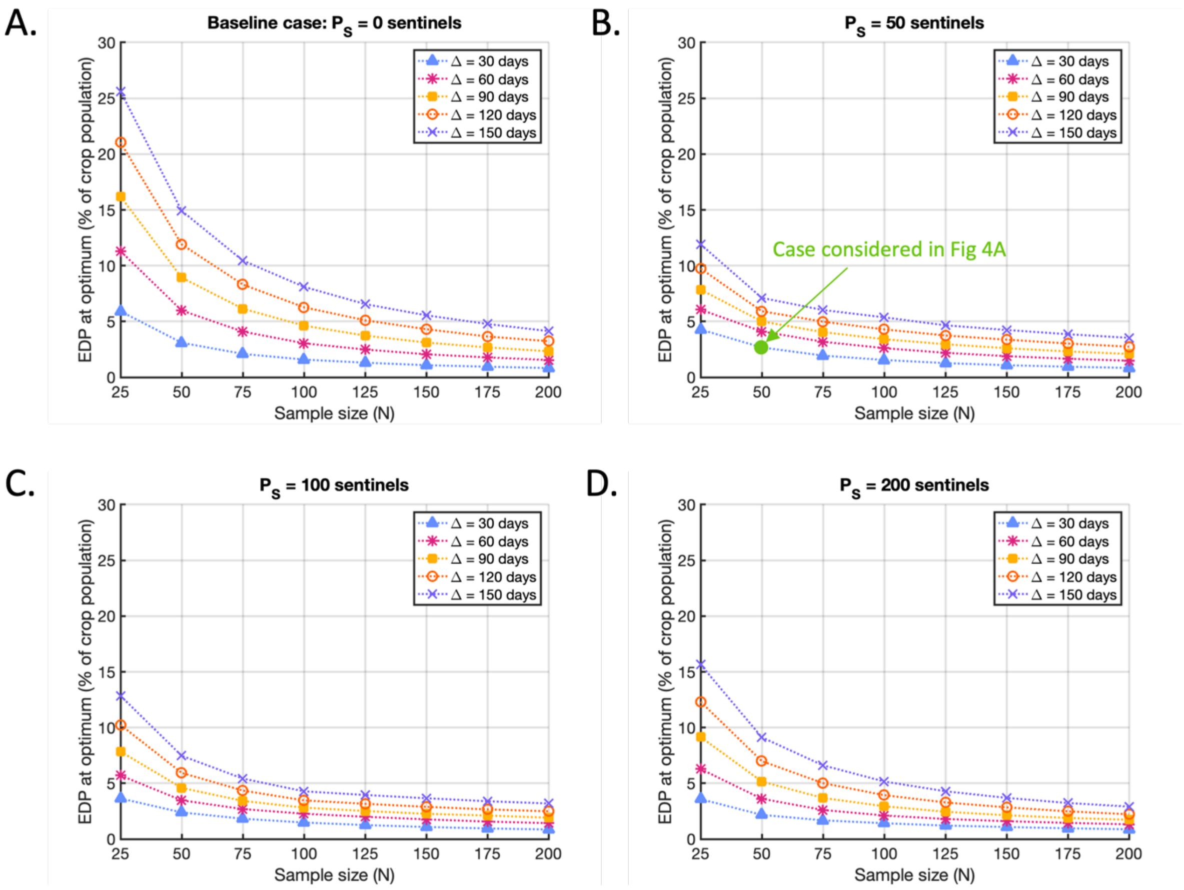


**Fig S16. Resultant EDPs in the baseline case and for the optimal strategies shown in Figs 4 and 5 of the main text.** A. The EDP in the baseline case ($P_S = 0$), where sample size ($N$) and sample interval ($\Delta$) vary as shown. Since the baseline EDP depends on $N$ and $\Delta$, relative changes in the EDP compared to this baseline for different values of $N$ and $\Delta$ (Figs 5ABC of the main text) are not a measure of the resultant EDP. B. The best achievable resultant EDP when $P_S = 50$, corresponding to the optimal strategies identified in Fig 4B in the main text. Green circle marks the case considered in Fig 4A in the main text ($P_S = 50, N = 50, \Delta = 30$ days). C. The analogous figure to B, but with $P_S =$ 100 sentinels added to the population and results corresponding to the strategies identified in Fig 4C. D. The analogous figure to B, but with $P_S = 200$ sentinels added to the population and results corresponding to the strategies identified in Fig 4D.

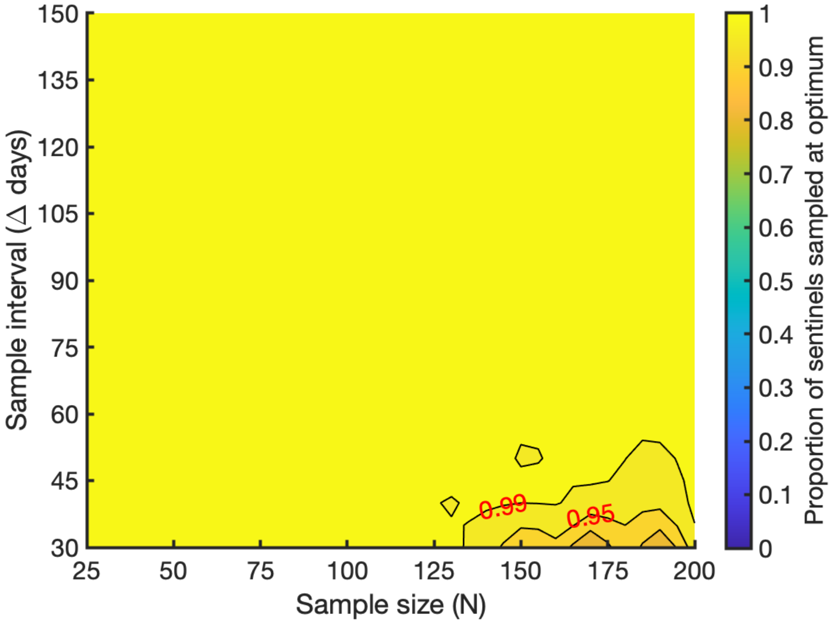


**Fig S17. The optimal proportion of available sentinels to include in the sample as the sample size ($N$) and sample interval (Δ) vary, for the system outlined in Section 3.3 and Fig 6 of the main text.** For each $(N, \Delta)$ pair, this proportion is computed as the optimal number of sentinels to include in the sample ($N_S^*$) divided by the maximum possible number of sentinels that could be included ($\min(P_S^*, N)$), where $N_S^*, P_S^*$ are obtained from the Bayesian optimisation algorithm.

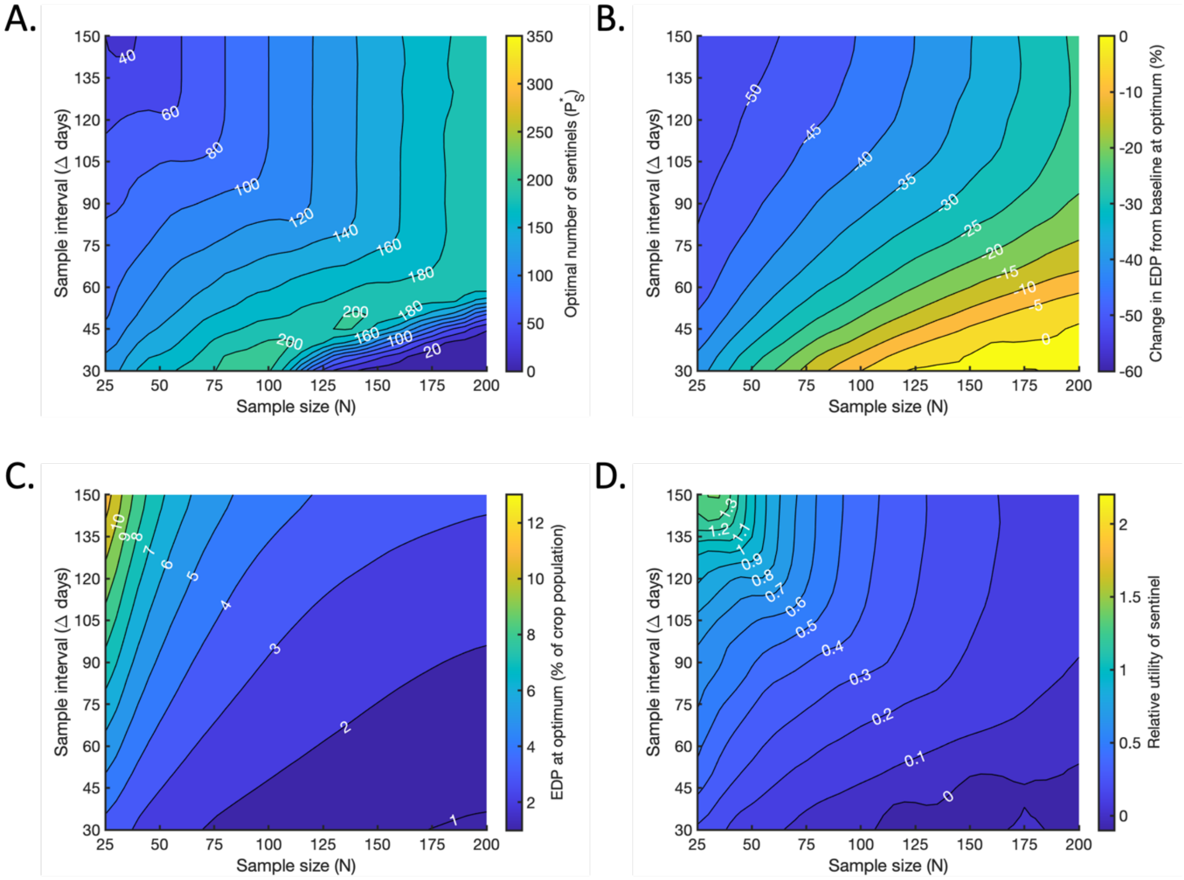


**Fig S18. Optimising the number of sentinels to include in the population and sample when $\boldsymbol{\Omega} = \mathbf{EDP} + (\mathbf{0.5} \times \mathbf{EDP_{sent}})$.** A. The optimal number $P_S^*$ of sentinel plants to include in the population, for which the maximal reduction in $\Omega$ compared to the baseline level is achieved (if $N_S$ is also chosen optimally). B. The percentage change in $\Omega$ compared to the baseline value at the optimum, achieved when $P_S = P_S^*$ and $N_S = N_S^*$. C. The resultant value of $\Omega$ at the optimum, expressed as a percentage of the total crop population. D. The relative utility of a single sentinel plant (the percentage reduction in $\Omega$ per sentinel) at the optimum.
